# Biological, freshwater, and marine drivers of age at maturity in wild Chinook Salmon

**DOI:** 10.64898/2025.12.13.694134

**Authors:** JL Gosselin, BP Sandford, CS O’Brien, ER Buhle

**Affiliations:** School of Aquatic and Fishery Sciences, University of Washington, Seattle, Washington, USA; Affiliate status: Northwest Fisheries Science Center, National Marine Fisheries Service, National Oceanic and Atmospheric Administration, Seattle, Washington, USA; Northwest Fisheries Science Center, National Marine Fisheries Service, National Oceanic and Atmospheric Administration, Pasco, Washington, USA; Mount Hood Environmental, Sandy, Oregon, USA

**Keywords:** age at maturation, carryover effects, freshwater and marine drivers, juvenile and adult Salmon, probit regression model

## Abstract

Understanding variation in age at maturity is important for endangered species recovery because older, larger adults contribute disproportionately to the next generation. Conditions in early life stages may have underappreciated impacts on age at maturity. Our study objective was to associate adult age of individually tagged wild, spring/summer Chinook Salmon (*Oncorhynchus tshawytscha*) outmigrating from the Snake River (Idaho and Washington, USA) in 1998–2020 with covariates measured during juvenile and subadult stages. We used a hierarchical Bayesian ordinal probit regression model to estimate statistical effects of juvenile body length, seasonal migration timing or river temperature, transported or in-river hydrosystem passage, river flow, and a large-scale ocean index. Results indicated notable carryover effects consistent with underlying biological mechanisms related to growth and development, in which shorter juvenile length and later seasonal migration timing were associated with older adults. These biological and behavioural factors were more important than riverine or marine environmental conditions examined. Our study suggests that managers and decision makers should consider carryover effects from the juvenile life stage on age structure in conjunction with survival.

## Introduction

Processes leading to older and more variable ages at maturity are important in helping to recover species listed under the US Endangered Species Act and to maintain their population sizes. Larger, older adult females will contribute more to reproductive output than smaller, younger adults that lay fewer eggs (Hixon et al. 2014, Ohlberger et al. 2020, Malick et al. 2023). Yet, in some systems, such as the Columbia River Basin in North America, adult age structure of Salmon and Steelhead (*Oncorhynchus* spp.) is often overshadowed in research and conservation management by investigations of abundance, recruitment, and survival (Ford 2022, NMFS 2022), even if declines in ages and sizes of adult Salmon have been occurring for decades (Ricker 1981, Ohlberger et al. 2018). Diversity in several Salmon life history traits, including age at maturity, is important for population viability (McElhany et al. 2000). Investigating the age structure of adult spawners in relation to biological and environmental covariates will contribute to our understanding of increased productivity and abundances, and thus inform strategies to increase population growth and reverse ongoing historical declines.

Declining age at maturity can indicate a fish population under stress (Trippel 1995). Influential anthropogenic factors are numerous and can include: overexploitation and selective removal of larger, older fish (Trippel 1995), hatchery production practices encouraging rapid juvenile growth (Vøllestad et al. 2004), timing of hatchery releases (Bilton et al. 1982), density-dependent effects of large hatchery releases, and compounding effects from a warming climate (Siegel et al. 2017, Connors et al. 2020). Despite these diverse causes, it is important to remember that the resulting adult age structure is a manifestation of both plastic responses (e.g., growth, smoltification, and developmental rates) and selection (or survival).

Variable age at maturity depends on factors associated with genetic, environmental, and physiological processes, as well as their interactions. Population genetics provides a baseline (Iwamoto et al. 1984, Larsen et al. 2019), and multiple factors influence physiological development in an ongoing process to maturation (Thorpe 2007, Spangenberg et al. 2015, Pauly 2022). Furthermore, environmental conditions can impact developmental thresholds (Day and Rowe 2002), which for salmonids can include ocean conditions in the year prior to maturation (Satterthwaite et al. 2019). Juvenile migration from freshwater habitats to the ocean is also a critical life stage for physiological development in anadromous fishes. High growth rate or large body size in juvenile salmonids is generally associated with younger age at maturity (Vøllestad et al. 2004, Shearer et al. 2006, Claiborne et al. 2011, Tattam et al. 2015). In addition to growth rate and length, seasonal migration timing and environmental conditions during downstream migration may be underappreciated drivers of age at maturity. Our study goal is to investigate these factors.

Passive integrated transponder (PIT) tags allow the tracking of conditions experienced by individual juvenile wild Chinook Salmon from the Snake River Basin as they migrate downstream through the Snake and Columbia rivers, to the Pacific Ocean and back to the river as adults. The timing of adults returning to the river to spawn can be tracked to determine the age at maturity (or more accurately the age of adult return because a fish may mature and die before being detected). The downstream migration of juveniles are hundreds of kilometres long and thus could have a significant influence on development. Historically, migration occurred under free-flowing hydrological conditions, but for many decades, the seasonal flow, temperature, and turbidity profiles in the Snake and Columbia rivers were and still are affected by impoundment of the Columbia River System (Figure 1). Downstream-migrating juvenile salmonids also have unique experiences as some of them are intercepted at dams on the Lower Snake River to be transported and released downstream of the dam closest to the ocean. Altogether, this case study provides a 25-year time series of environmental conditions and fish experiences in a highly altered and regulated river system. Examining environmental, biological, and behavioral covariates in the juvenile stage could help to better understand drivers associated with age structure of returning adults.

**Figure 1.**
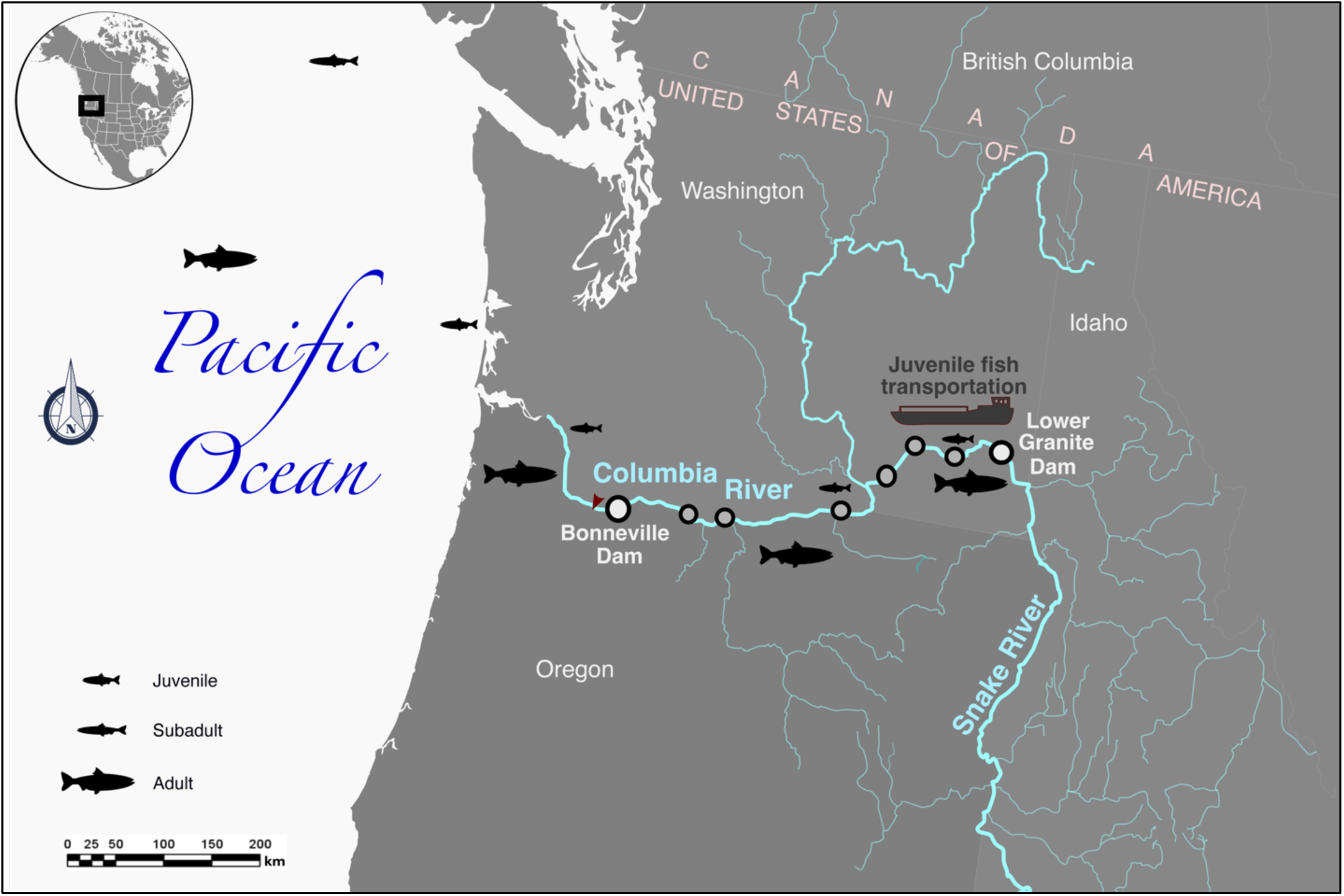
Map of the Snake River, Columbia River and their tributaries, Pacific Northwest, USA. Juvenile wild spring/summer Chinook Salmon migrate in-river or are transported in barges downstream through the hydrosystem to the ocean, where they reside for 1–3 years before maturing and returning to spawn.

Our study objective is to examine patterns in the adult age structure of wild spring/summer Snake River Chinook Salmon and their relationships with potential biological and behavioural covariates (fork length, age and location at tagging and release, origin, and migration timing), freshwater environmental covariates (in-river vs transported hydrosystem passage experience, river temperature and flow), and a marine covariate (North Pacific Gyre Oscillation index). Identifying and quantifying covariates associated with adult age structure, and consequently fecundity, would provide information applicable to the recovery of endangered species and maintenance of viable populations for at-risk populations.

## Methods

### Study system

The Snake and Columbia rivers (Pacific Northwest, USA) were once free-flowing, but the construction of multiple dams from the 1930s through the 1970s have transformed them into largely a series of reservoirs. In the current study, we examined wild spring/summer Chinook Salmon that migrate through eight major dams (hereafter the “Columbia River System” or “hydrosystem”; Figure 1). As part of hydrosystem operations, the Juvenile Fish Transportation Program (US Army Corps of Engineers) increases the survival of downstream-migrating juveniles from approximately 40-60% to nearly 100% by using barges outfitted with a filtration system to transport juveniles collected at the three furthest upstream dams and release them below the furthest downstream dam. Despite these immediate increases in survival, delayed mortality expressed in the post-hydrosystem environment may outweigh the benefits in some years (Gosselin et al. 2018, Gosselin et al. 2021a).

### Study species and covariates

We examined wild, spring/summer Chinook Salmon from the Snake River Basin that were tagged with PIT tags (Columbia Basin PIT Tag Information System [PTAGIS], Pacific States Marine Fisheries Commission, Portland, Oregon, USA; www.ptagis.org/). Juveniles were tagged and released at Lower Granite Dam (LGR; river kilometre [rkm] 695, with rkm 0 at the mouth of the Columbia River) or further upstream. Our data set included fish that were detected as juveniles at LGR from 1998 to 2020 and subsequently detected as adults at Bonneville Dam (BON; rkm 234) or any further upstream dam (Figure 1).

Many Chinook Salmon were designated at the time of tagging as an unknown run, so we filtered the dataset for wild spring/summer Chinook Salmon as opposed to counterparts that were hatchery-reared or fall-run. We included juveniles with fork lengths 45 to 155 mm and passage detections at LGR on day of year (DOY) 80 to 160, based on our knowledge on different runs and life stages of Chinook Salmon and historical data. We excluded any adult returns with PIT-tag detection patterns indicating ages that were 2 y (or 0-ocean; mini-jacks) and > 6 y (or > 4-ocean), with the assumption that all juveniles were 2 y of age when entering the ocean. Two fish aged 6 y were assigned to the same adult-age category as fish aged 5 y (or 3-ocean) (Table S1 in Suppl. Mat.). We excluded adults that passed BON in October–December because these were likely fall-run Chinook Salmon, which is a timing in alignment with other studies (Crozier et al. 2020, Coykendall et al. 2022). In addition, for juveniles that were tagged and released at LGR (i.e., run-at-large Chinook Salmon of unknown origin), we excluded adults that passed BON in August-December because they were potentially fall-run based on study information and comments noted in PTAGIS. In total, we analyzed 8,019 wild spring/summer Chinook Salmon adult returns (for details on sample sizes, see Tables S1, S2, and S3 in Suppl. Mat.).

We investigated biological/behavioural, freshwater, and marine covariates hypothesized to affect the age of adult returns (Table 1). The biological/behavioural covariates included juvenile fork length at tagging, DOY of LGR passage, and origin, which was determined based on the Major Population Group (MPG; McClure et al. 2005) and tag release group by juvenile life stage (subyearling parr vs yearling smolt) and location. The covariates representative of freshwater experiences included hydrosystem passage type (in-river vs transported) as well as river flow and temperature at LGR at the time of passage. Because DOY of passage was highly correlated with river temperature (*r* = 0.76), only one of these two covariates was included in the model at one time. The marine index examined was the North Pacific Gyre Oscillation (NPGO) index (Di Lorenzo et al. 2008), averaged over the fall-winter months (October–March) across 3 years, starting with October in the year of smolt ocean entry. Other marine and climate indices, such as the Pacific Decadal Oscillation index and the Aleutian Low Pressure Index, were examined in preliminary analyses. However, only one index was used because of the changes in the correlations among these large-scale indices (Litzow et al. 2020, Gosselin et al. 2021b). As well, the NPGO was chosen for its association with Salmon responses and other species in the ecosystem of the Pacific Ocean (Kilduff et al. 2015, Malick et al. 2017, Couture 2024). All continuous covariates were standardized to mean = 0 and standard deviation = 1. Categorical covariates were passage type (in-river or transported), release life stage and location (parr above LGR, parr at LGR, and smolt at LGR), and origin (MPG and tag and release location).

**Table 1.** Potential covariates of adult age, data sources, and hypothesized mechanisms.

| Covariate name | Description | Data source | Hypothesized mechanisms |
| --- | --- | --- | --- |
| **Origin** | Origin of Chinook Salmon grouped by Major Population Group and sampling location: Lower Snake (i.e., Lower Granite Dam [LGR] samples), South Fork Salmon, Middle Fork Salmon, Lower Salmon, Upper Salmon, Clearwater, Grande Ronde/Imnaha. | www.ptagis.org/ | Differences in genetics and rearing conditions lead to differences among their physical and physiological states as juveniles and consequently their age at maturity. |
| **Tagging life stage & location** | Release group of tagged juveniles by life stage and above/at LGR: parr above LGR, smolt above LGR, or smolt at LGR. | www.ptagis.org/ | Tagging effects result in differential culling and/or differential growth and development at parr and smolt life stages, and consequently result in different ages at maturity. There may be population effects as well due to different populations sampled above vs at LGR. |
| **Length** | Fork length (mm) from snout to fork in the tail at time of tagging and release; limited to range (45–155 mm), as expected for wild Snake River Chinook Salmon parr and smolts. | www.ptagis.org/ | Juveniles with slower growth rates and smaller sizes mature at older ages. |
| **Length × Tagging life stage & location** | Interaction effect | Same as above | The length effect on age at maturity differ for parr vs smolts, and for different populations sampled above vs at LGR. |
| **DOY** | Day of year (DOY) of juvenile passage at LGR, limited to DOY 80–160. | www.ptagis.org/ | Later-migrating fish are on a slower growth and development schedule, and thus mature at a later age. |
| **Passage type** | Presence/absence (1/0) of juvenile transportation: juveniles intercepted through the bypass system at LGR are put on barges for transportation to below the most downstream dam (Bonneville Dam) or returned to the river for in-river passage. | www.ptagis.org/;<br>www.cbr.washington.edu/dart/metadata/pit#transport | Transported juveniles experience disrupted and stressful passage through the hydrosystem and minimize their mortality risk by maturing earlier because of the perceived unfavourable conditions.<br>Alternatively, smaller, older-maturing fish that experience juvenile transportation subsequently undergo size-selective mortality. These potential older adults do not survive to return to the river. |
| **Flow** | 7-d mean of river flow (kcfs) at LGR; mean is right-aligned to fish passage date | Water Quality Monitoring data from USACE; accessed at www.cbr.washington.edu/dart/ | Juveniles that experience higher flow are less stressed and mature at an older age because of the perceived favourable conditions.<br>Alternatively, smaller, older-maturing fish that experience high flow have higher ocean survival. These older adults survive to return to the river. |
| **Temperature** | 7-d river temperature (°C) at LGR; mean is right-aligned to fish passage date | Water Quality Monitoring data from USACE; accessed at www.cbr.washington.edu/dart/ | Juveniles that experience higher temperatures minimize their mortality risk and mature at a younger age because of the perceived unfavourable conditions.<br>Alternatively, smaller, older-maturing fish that experience high temperatures have lower ocean survival. These older adults do not survive to return to the river. |
| **NPGO** | North Pacific Gyre Oscillation (NPGO) index, averaged for the fall-winter months (ONDJFM) prior to adult return | Di Lorenzo et al. (2008); http://www.o3d.org/npgo/ | Juveniles and subadults decide to stay in favourable ocean conditions (positive values of NPGO) to grow to a larger size and increase fecundity potential, thus mature at an older age.<br>Alternatively, small-sized fish that mature at older ages have higher ocean survival in favourable ocean conditions. These older adults survive to return to the river. |

### Model

To quantify the relationships between age of adult returns and biological/behavioural, freshwater, and marine covariates, we used a hierarchical Bayesian ordinal probit regression model (Gelman and Hill 2007, Jackman 2009, Bürkner and Vuorre 2019). This model is formulated as a multilevel regression with covariates **x** and a latent, unobserved continuous response 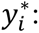

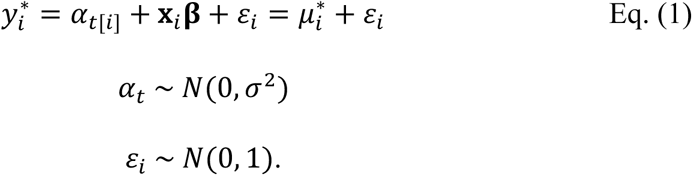

Here **x***_i_* is the length-*K* row vector of covariates for individual *i*, **β** is the vector of regression coefficients, and *α_t_*_[*i*]_is the group-level intercept (or “random effect”) corresponding to the year *t* in which individual *i* entered the ocean. The year-level intercepts follow a normal hyper-distribution with standard deviation (SD) *σ*. The residuals *ε_i_* have standard deviation fixed at 1 for identifiability, as described below.

The observed ordered categorical response *y* ∈ [1, 2, 3] for ages 1-ocean, 2-ocean, and ≥ 3-ocean (or total ages 3, 4 and ≥ 5 y), respectively, is related to the underlying continuous 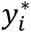 by censoring, such that:

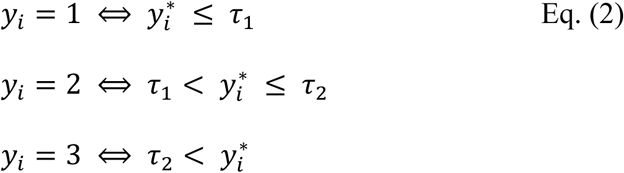

where ***τ*** = [*τ*_1_, *τ*_2_] is a vector of cutpoints sorted in ascending order. This implies that the probabilities of each categorical outcome are:

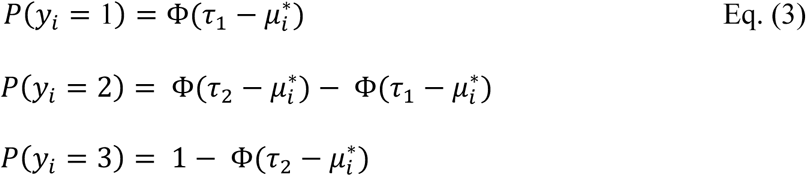

where Φ(·) is the standard normal cumulative distribution function (see a graphical depiction of the probability density function in Figure 3). Some constraints are required for parameter identifiability in this model; we fix the global intercept of *y*^∗^at zero (so **x***_i_* does not include a constant term) and the residual SD at 1, ensuring that the regression slopes **β** and the cutpoints **τ** are identified (Jackman 2009). The group-level intercepts *α_t_* shift 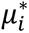 up or down for each outmigration year *t* around the global intercept of zero, which is equivalent to shifting all the cutpoints by −*α_t_*.

We fit the model using Hamiltonian Monte Carlo sampling (Monnahan et al. 2017) as implemented in the Stan platform via the `brm` function from the `brms` package (v.2.20.4; Bürkner 2017, 2018). The regression coefficients **β** and the cutpoints **τ** were given weakly informative priors (Lemoine 2019) with diffuse normal distributions, *N*(0,5), and the hyper-SD *σ* was given a weakly informative prior of Student’s *t*(3,0, 2.5). We simulated 4,000 draws from the posterior distribution in each of 3 randomly initiated chains, discarded the first 1,000 draws as warmup, and thus saved a total of 9,000 draws. To assess convergence, we inspected traceplots and density plots, required potential scale reduction factor *R̂*^F^< 1.02, and had no divergent transitions (Gelman et al. 2014). Additionally, the `loo` function from the `brms` package was used for model selection of candidate submodels based on approximate leave-one-out cross-validation to assess the relative importance of biological/behavioural, freshwater, and marine covariates (Vehtari et al. 2017).

## Results

In smolt migration years 1998–2020, most of the returning adults from the PIT-tagged juvenile wild spring/summer Snake River Chinook Salmon were age 2-ocean (4 years total age; Figure 2). Overall, the proportion of 3-ocean adults declined over time (Figure 2b). Variation around this trend did not appear to be related to adult counts. For example, the lowest proportions of 1-ocean adults were from smolt migration year 2000, which had a high count, and from 2003–05, which had relatively low counts. There were also more 3-ocean than 2-ocean adults in 2000. By contrast, migration years 2015–2016 had low proportions of 3-ocean adults and were among the cohorts with the fewest adult returns (Figure S1 in Suppl. Mat.). Migration years 2008–2009 had typical age distributions but relatively high adult returns.

**Figure 2.**
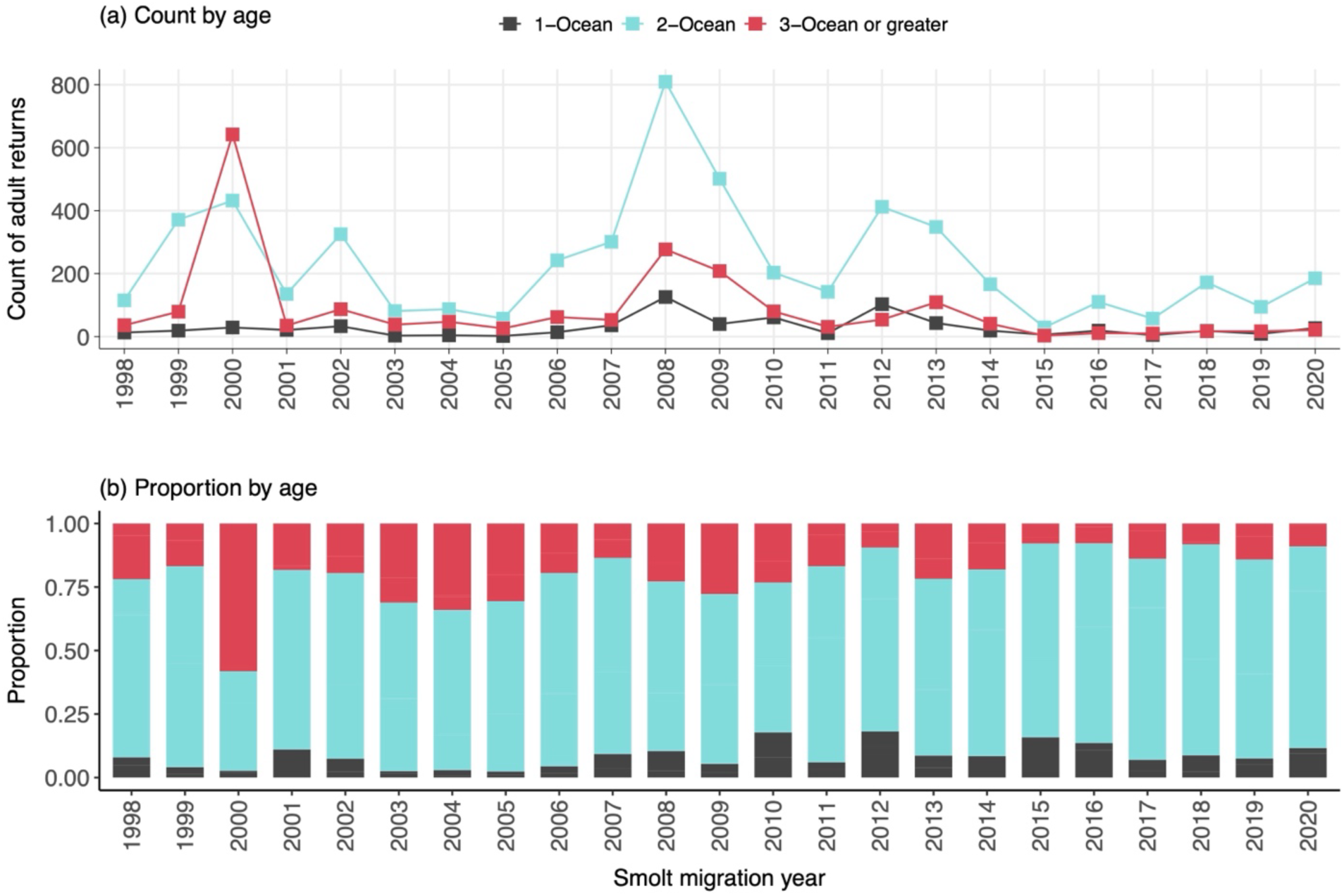
Annual counts (a) and proportions (b) of wild Snake River spring/summer Chinook Salmon by adult age

**Figure 3.**
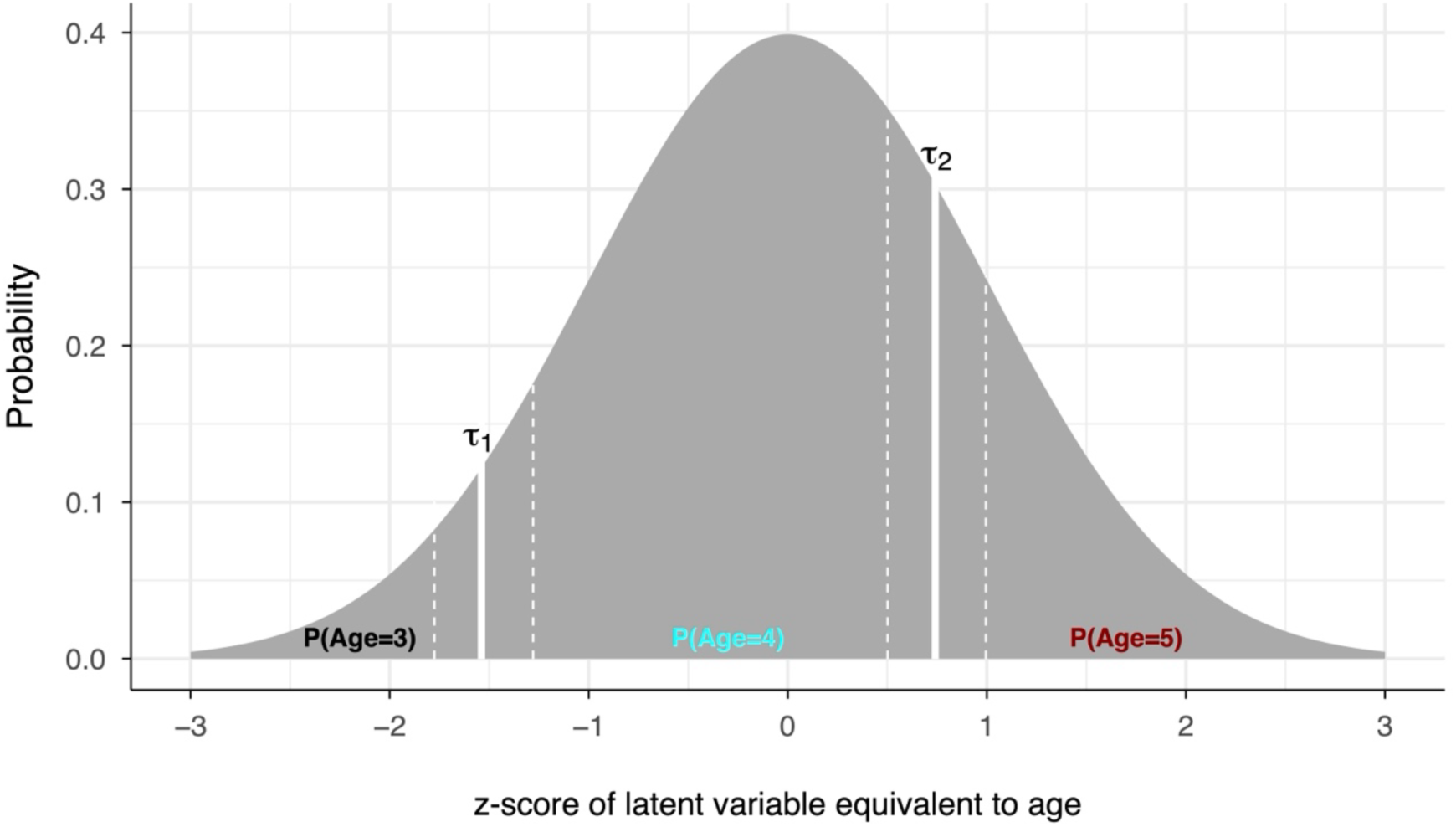
Probability density function for a hypothetical ordinal probit regression model. The probability of the latent variable for each age is the relevant area under the curve, determined by the cutpoints τ. A covariate with positive effect β would shift τ left, capturing a larger proportion of older individuals; conversely, a negative β would shift τ right, capturing a larger proportion of younger individuals (Eq. 3).

Outmigrating juvenile and subadult Chinook Salmon were transported downstream at rates ranging from 21 to 93% (Figure S1 in Suppl. Mat.). Juvenile fork length was 46–153 mm (Figure S5 in Suppl. Mat.). Smolts passed LGR from 25 March–9 June (Figure S6 in Suppl. Mat.), when water temperature was 6.0–14.4 °C (Figure S7 in Suppl. Mat.) and flow was 29–189 kcfs (Figure S8 in Suppl. Mat.). The NPGO index generally decreased over time, with peaks in 2000 and in 2007 (Figure S9 in Suppl. Mat.).

Assuming independence among covariates, the full model indicates that the fish with the highest probability of returning at older ages are smaller juveniles tagged as smolts (rather than parr) above LGR and migrating later in the season, passing through the hydrosystem in-river at high flows, and entering a productive ocean with a high NPGO index (Figure 4). Thus, indirect effects on age at maturity may carry over from the freshwater juvenile life stage to the adult life stage returning from the ocean. Adult age distributions also varied by origin; younger adults originated from SF Salmon and MF Salmon, while the Lower Snake, Clearwater, and Upper Salmon tended to produce older adults.

**Figure 4.**
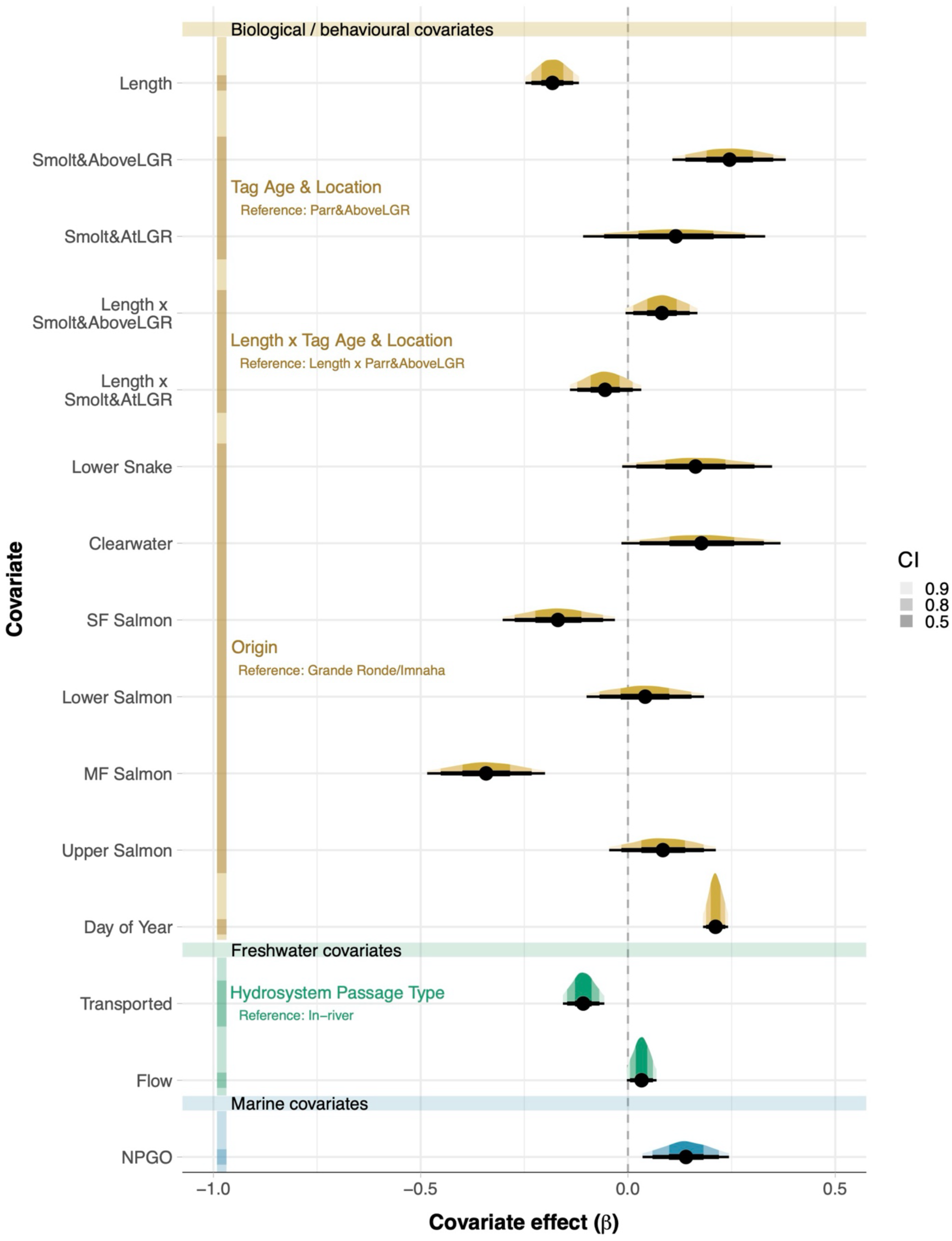
Estimated regression coefficients (β) for effects of covariates on adult age of wild Snake River spring/summer Chinook Salmon.

The full model with DOY of juvenile passage at LGR performed the best, based on approximate leave-one-out cross-validation. Estimated effects were qualitatively similar when DOY (Table S4 in Suppl. Mat.) was replaced with river temperature (Table S5 in Suppl. Mat.), but the latter model did not fit the data as well. A model with only biological/behavioural covariates gave a more parsimonious fit than either the full model with river temperature in place of DOY, the model with only freshwater covariates, or the model with only NPGO (Tables S5–S10 in Suppl. Mat).

Conditional effects plots show the relative effect of each covariate, with all others held at their sample means (Figure 5). The most notable effects were juvenile body size and outmigration timing. The estimated probability of younger maturation increased with juvenile fork length (Figure 5a). For example, a 150-mm juvenile is approximately twice as likely to return as a 1-ocean adult than is a 110-mm juvenile. Conversely, a 135-mm smolt is half as likely to return as a 3-ocean adult than is a 100-mm smolt. A fork length of 130 mm compared to 120 mm decreases the median probability of returning at ocean age 3 from 0.135 (90% CI 0.092–0.191) to 0.111 (90% CI 0.069–0.167). This translates into 24 fewer 3-ocean adults out of every 1000. For juveniles tagged and released above LGR, the length effect was greater in smolts than in parr (Figure 5c,e). The origin groupings, in order of youngest to oldest median adult ages, are MF Salmon, SF Salmon, Grande Ronde/Imnaha, Lower Salmon, Upper Salmon, Lower Snake, and Clearwater (Figure 5f). The effect of LGR passage timing was also notable, with the median probability of 3-ocean adult return increasing from 0.066 (90% CI 0.043–0.093) to 0.365 (90% CI 0.289–0.446) over the migration season (Figure 5g). Across a range of increasingly favourable ocean conditions, with NPGO values of –2.07 to 1.92, the median probability of 3-ocean adult return increased from 0.106 (90% CI of 0.061–0.160) to 0.231 (90% CI of 0.159-0.311; Figure 5j).

**Figure 5.**
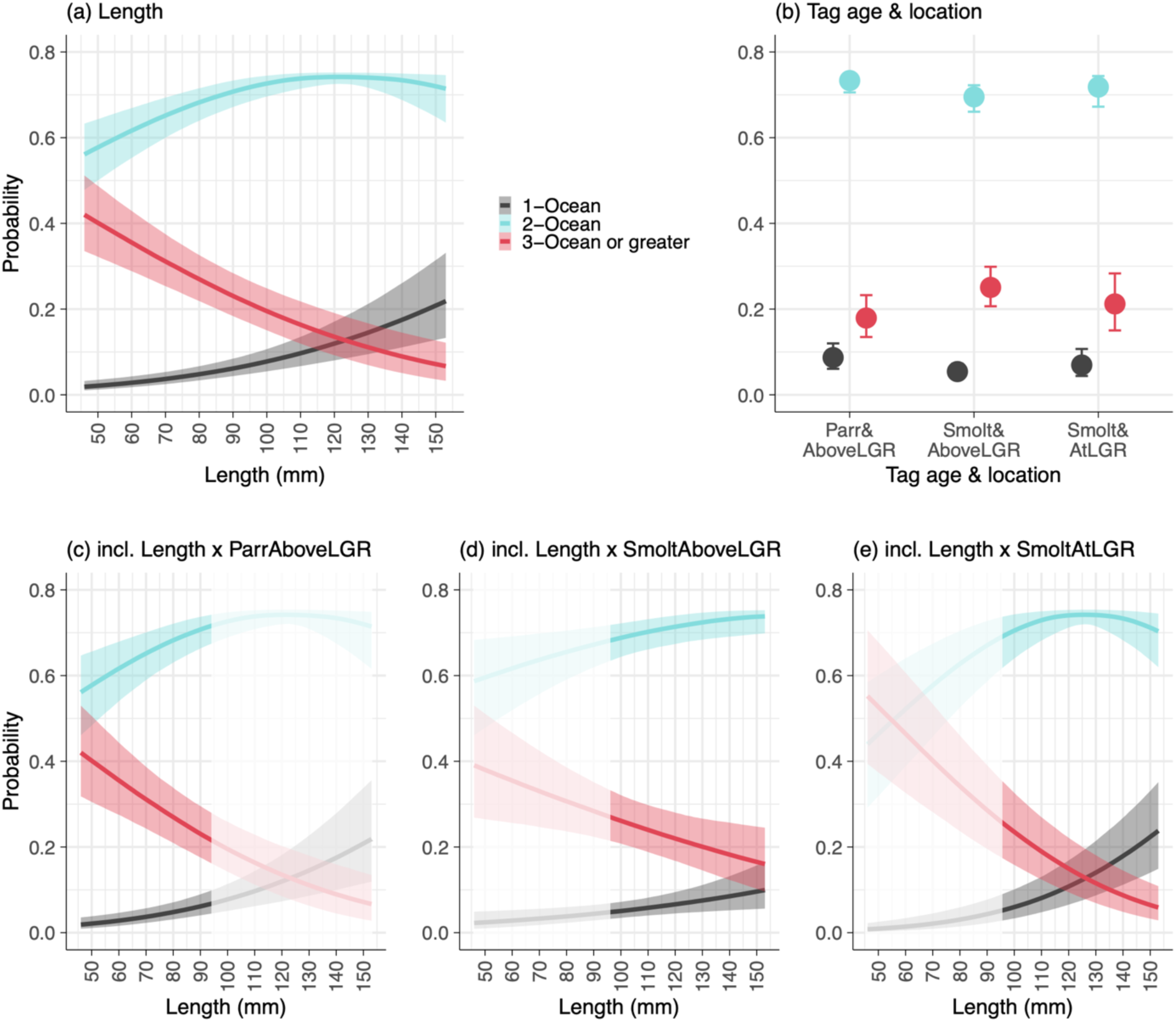
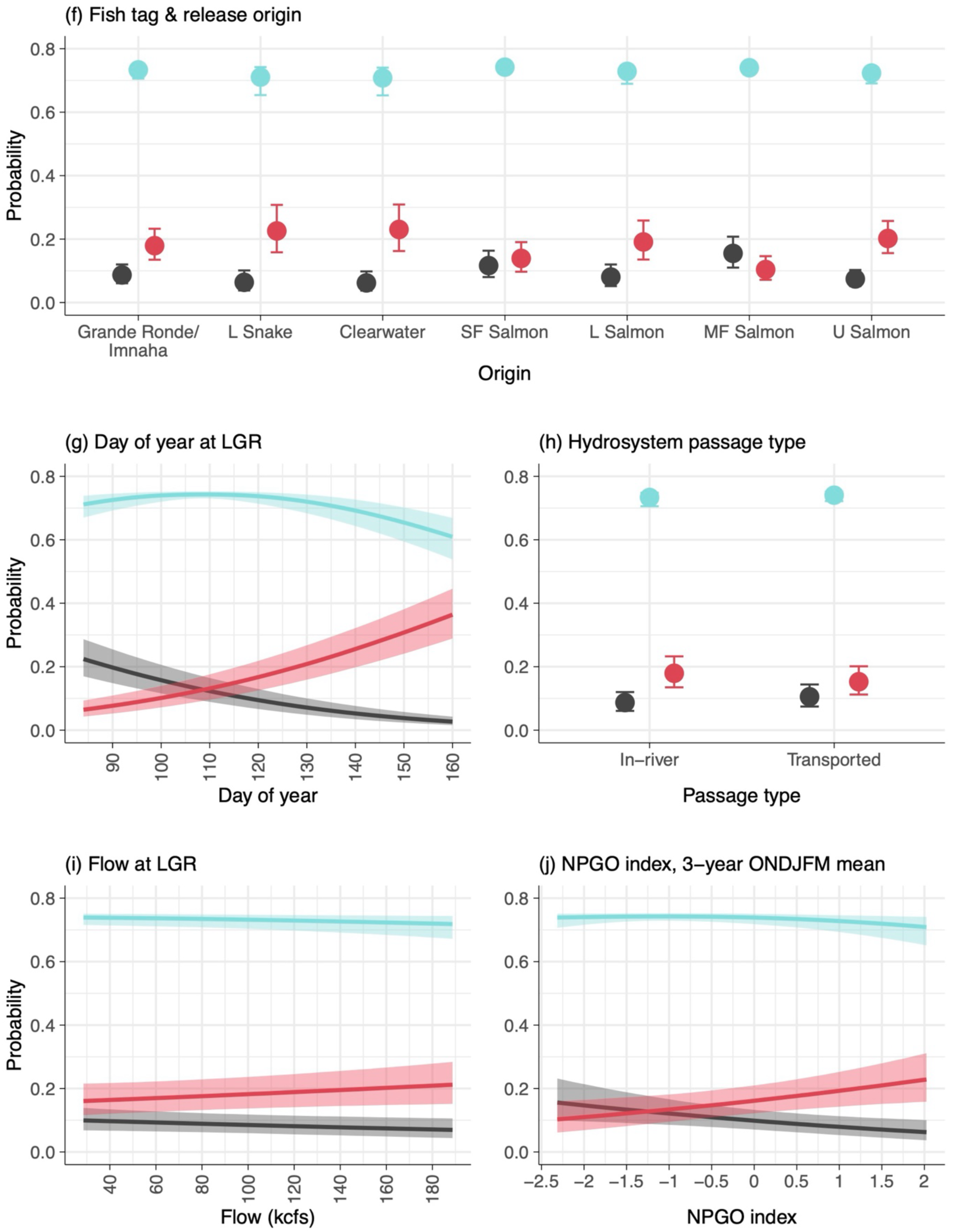
Conditional effects of individual covariates on age-at-return probabilities of adult Chinook Salmon, with all other covariates held at their sample means. Lines and dots represent medians, and shaded areas and error bars represent 90% credible intervals. Faded sections of 4c, 4d, and 4e represent areas of extrapolation beyond the observed values.

Predicted adult age distributions in each smolt migration year were close to the observations, even without including the annual random effects (Figure 6). Nevertheless, the annual random effects can still be relatively large (Table S11 in Suppl. Mat.); for example, the positive random effect in the year 2000 shifted the cutpoint further left to capture the greater proportion of 3-ocean than 2-ocean adults that our covariates did not explain.

**Figure 6.**
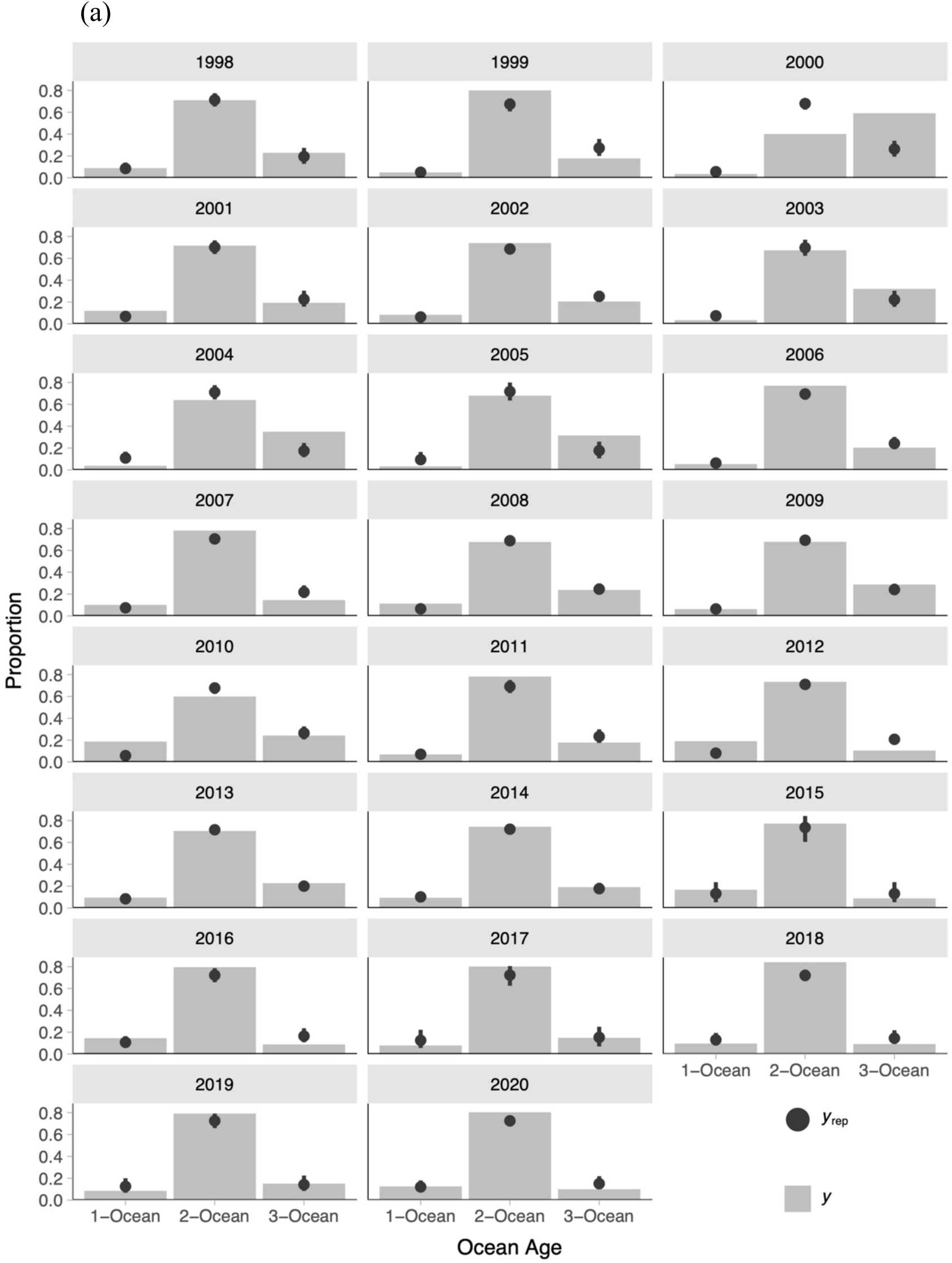
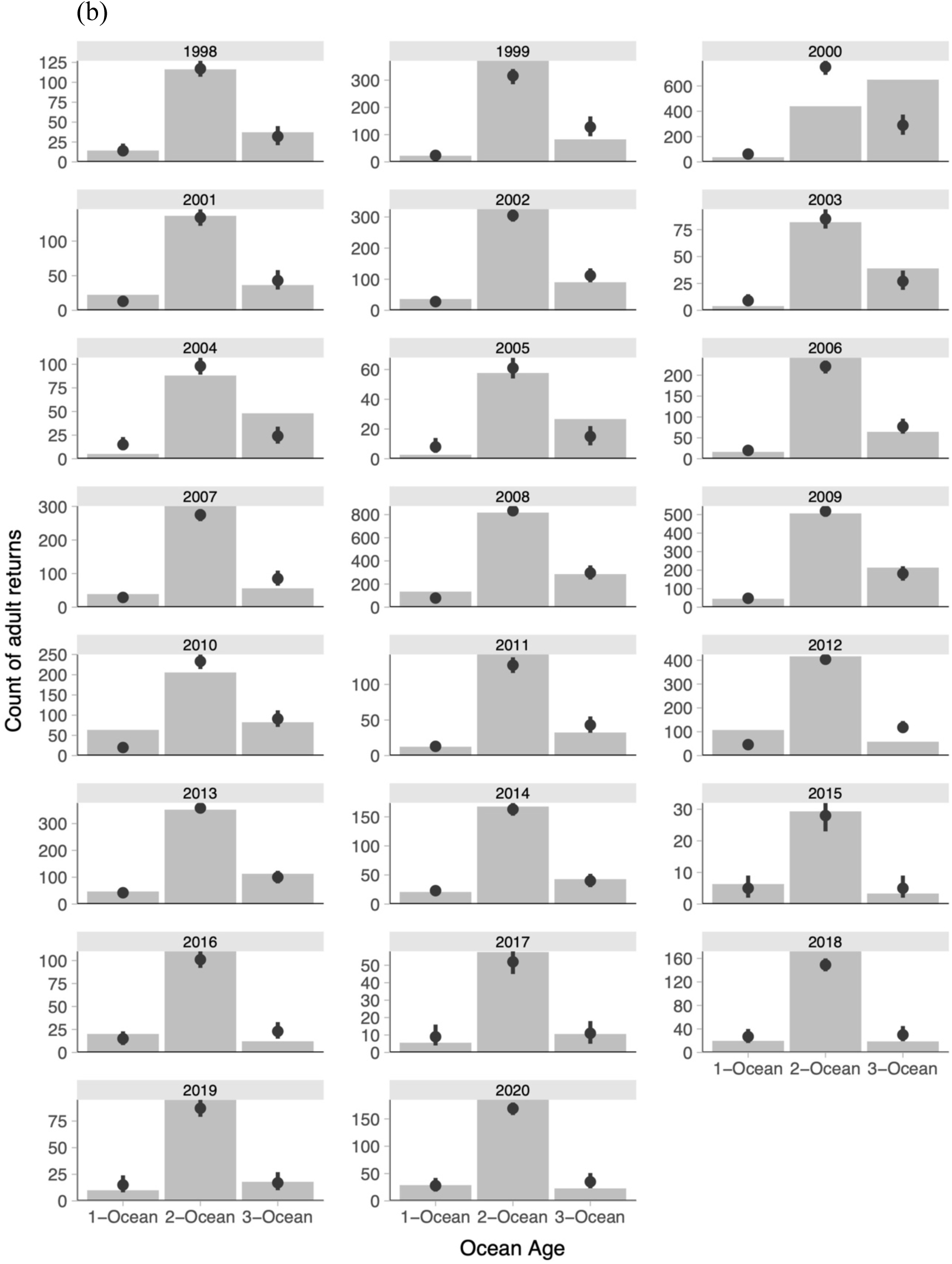
Predicted and observed adult age distributions of wild Snake River spring/summer Chinook Salmon by juvenile outmigration year, shown as relative frequencies or proportions (a) and as raw frequencies or counts (b). Bars represent observations, and points and error bars represent posterior medians and 90% credible intervals, respectively. Predictions correspond to the hyper-mean, i.e. they do not include annual random effects. Note that credible intervals in some cases are too narrow to be visible.

## Discussion

Several covariates associated with the freshwater juvenile life stage influenced the age at maturity of wild spring/summer Chinook Salmon from the Snake River Basin. All else being equal, adults were more likely to return at older ocean ages if they were smaller-sized juveniles that outmigrated through the hydrosystem, later in the season, in higher river flows. In addition, favourable ocean conditions predicted greater age at maturity. These patterns may seem counterintuitive, given that older adults were associated with slower juvenile growth but hypothetically more rapid subadult growth at sea. Interpreting these results requires a lens that integrates effects across life stages, which we discuss further below. Moreover, adult age distributions varied with the juvenile life stage (parr or smolts) and location of tagging (at or upstream of LGR) as well as the subbasin of origin. Overall, the biological/behavioral covariates (juvenile size and migration timing) had stronger effects on the adult age distribution than the environmental drivers (in-river passage, river flow, and NPGO). In the following discussion we interpret our findings in the context of drivers of age and survival, and of the hydrosystem, hatcheries, harvest, and habitat, collectively known as the four H’s that have impacted Salmon populations in the Columbia River System.

### Drivers of age at maturity and survival

Interestingly, some covariates associated with greater adult age have previously been linked to lower survival (e.g., shorter juvenile lengths and later outmigration in the season; Sogard 1997, Scheuerell et al. 2009, Gosselin et al. 2021a), while others have been linked to higher survival (e.g., productive ocean; Kilduff et al. 2015). These seemingly paradoxical relationships can be explained by an interaction of juvenile-to-adult carryover effects with size-selective mortality. If smaller, later-migrating smolts are less likely to survive and also more likely to spend more time at sea before maturing, then favourable ocean conditions may ameliorate their survival disadvantage and allow more of them to return and thus be included in our sample. This also suggests potential tradeoffs in population productivity, mediated by countervailing effects of factors such as juvenile size on marine survival on the one hand, and age at maturity (a proxy for size-specific female fecundity) on the other. Additional covariates that are believed to increase survival, namely flow and in-river hydrosystem passage, were positively related to age at maturity but these effects were relatively small.

### The four H’s

Regarding the hydrosystem, we found that older adults were associated with the more natural conditions of in-river passage and higher flow. Our analysis cannot distinguish between the alternative hypothesized mechanisms underlying these two effects (Table 1), i.e. whether these drivers directly increased age at maturation, improved the relative marine survival of smaller and older-maturing fish, or both. In any case, passage type and flow had relatively small effects compared to the biological/behavioural covariates over their observed ranges. Based on leave-one-out cross-validation, our models supported an effect of migration timing more strongly than an effect of river temperature. This is noteworthy, considering the multicollinearity between the two covariates. Altogether, these results suggest that genetic, developmental, and physiological processes throughout the juvenile life stages influence adult age more strongly than environmental conditions experienced during a relatively brief period of the life cycle.

The effect of juvenile length in our analysis is consistent with previous studies showing that faster juvenile growth, larger body size, and early outmigration timing are associated with early sexual maturation in hatchery Salmon (Larsen et al. 2006, Shearer et al. 2006, Bosch et al. 2023) and to a lesser extent wild Salmon (Siegel et al. 2018). These results suggest that broodstock selection and rearing conditions may help determine the maturation schedule of hatchery-origin fish. Based on our study of wild Chinook Salmon, a relatively modest decrease in smolt length of 10 mm could increase the number of older adults by roughly 24 fish per thousand. Any practical application of this inference would of course be subject to species-, population-, and broodstock-specific variation, since genetics and culture practices may dominate the effect of size at release.

Adult Chinook Salmon from the Clearwater River had an older age distribution than other origin groups in our study. This was unexpected because anadromous fish passage in this subbasin has been blocked or reduced by hydroelectric dams during multiple periods (1927–1940s near the mouth of the Clearwater River, late 1920s–1963 in the SF Clearwater River, and 1970–present in the NF Clearwater River), and as a result hatchery production of Salmon and Steelhead has been used for mitigation and restoration (Lindland and Bowler 1988, Chewning and Starr 2022). Hatchery-origin and hatchery-influenced wild populations often show a shift toward younger adult age (Riddell et al. 2024), but in this case we found the opposite. One possible explanation is that broodstock management practices in the Clearwater subbasin may have selected for older spawners. Alternatively, juveniles may experience slower growth due to relatively cold temperatures in the Clearwater River, but we already account for a length effect in our model. This example illustrates how population- and location-specific variation may dominate general patterns associated with rearing type.

Size-selective fisheries can alter the age structure of fish populations (Allendorf and Hard 2009), often by targeting individuals above a size threshold and thus driving size and age downward (Sharpe and Hendry 2009), although this is not always the case (Kendall et al. 2014). These dynamics involve complex interactions between fishery mortality and both genetic and plastic responses; for example, fishing may reduce abundance and release density-dependent growth, leading to earlier maturation. In general, however, evidence from modeling, laboratory experiments and field observations indicates that commercial fisheries typically shift populations toward smaller and younger demographic composition (Sharpe and Hendry (2009). Such shifts have implications for population viability, ecosystems and fisheries (Oke et al. 2020). In the case of recreational and commercial Chinook Salmon fisheries in the Columbia River Basin, Kendall and Kostow (2016) estimated that length- and age-based standardized selection differentials were relatively small. If this result continues to hold with more recent data, it may suggest that size-selective harvest does not adequately explain the trend of declining 3-ocean adult proportions that we observed.

Habitat restoration for Salmon populations often aims to increase carrying capacity and thus abundance (Lorenz et al. 2013, Hinrichsen and Paulsen 2020, See et al. 2021, Beechie et al. 2023). This may involve reconnecting floodplains, adding complexity to simplified stream channels, and restoring impaired flow and thermal refugia (NMFS 2022) in order to relieve density dependence, increase food resources, and improve juvenile growth (Ebersole et al. 2006, Bond et al. 2019, Luis et al. 2024). Growth rates may either increase or decrease depending on geographic location, elevation, and changes in thermal regime associated with climatic warming (Crozier et al. 2008, Beer and Anderson 2011, 2013), as well as alternative juvenile migration strategies such as natal or downstream rearing (Copeland et al. 2014). This diverse suite of habitat-mediated influences on growth can in turn affect the maturation schedule. Habitat restoration has been shown to increase the relative abundance of older age classes in some cases, e.g. brown trout in Spain, but this topic is understudied (Antón et al. 2011, Louhi et al. 2016).

We propose that the four H’s do not have equal influence on the adult age structure of Snake River spring/summer Chinook Salmon. Hatchery operations can most directly impact juvenile growth and size and consequently age at maturity. By contrast, hydrosystem conditions such as flow and passage route have detectable but relatively modest effects on adult age composition. Data and knowledge gaps are greatest with respect to harvest and habitat influences on age structure. Considering the severe decline in abundance of all species of wild Salmon in the Columbia River Basin from a total of 11–15 million in the late 1800s to 0.1–0.3 million in the late 1900s (Gresh et al. 2000, Lackey 2003), accompanying decreases in age and size also likely occurred. Effects of habitat restoration on age structure are challenging to study due to the decadal time scales and the complexity of direct and indirect effects. Long-term monitoring and assessment is still needed to understand how population demographic composition responds to conservation actions.

### Future research and monitoring

Adult age structure, as well as smolt and adult body size, have implications for the recovery and viability of salmonid populations and should be monitored and included in status evaluations (McElhany et al. 2000, Ford 2022, NMFS 2022). Declining adult age and size may lead to lower fecundity (Hixon et al. 2014, Ohlberger et al. 2020, Malick et al. 2023), inability to reach spawning grounds (Hinch and Rand 1998, Hinch et al. 2021, Twardek et al. 2022, Wilson et al. 2022, Birnie-Gauvin et al. 2023), and diminished capacity to dig redds deep enough to avoid scour (DeVries 1997, Adelfio et al. 2019, Parasiewicz et al. 2019). Greater variation in age structure with more representation of older adults can also provide a temporal buffer against the risk of unfavourable environmental conditions (Hixon et al. 2014). Recent status reviews and viability assessments include annual age and size composition for some populations, but these data remain underrepresented relative to metrics such as total adult abundance. We therefore emphasize the recommendation for more regular sampling and reporting. Moreover, it is important that age-composition data be reported as raw frequencies (counts) rather than relative frequencies (proportions) whenever possible, as the sample size conveys information about precision that is crucial for statistical estimation of underlying demographic parameters.

Further research could address potential biases in our methods and refine our results. One limitation of our study is that PIT-tagged fish may not be a representative sample from the population size distribution. For example, Coykendall et al. (2022) found that parentage-based tagging gave more accurate abundance estimates than PIT-tagging. Age-composition sampling based on noninvasive methods such as fish scales (Johnson et al. 2012, Wright et al. 2015, Quist and Isermann 2017) would be a useful point of comparison. Long-term monitoring of age structure in adult scales (Todd et al. 2014) and growth in juvenile scales (Gosselin et al. 2024) could be informative and relatively easy to implement. Some populations have undergone declines in both size and age structure, while others have had no change in age structure and only declining size-at-age (Morita and Fukuwaka 2007, Kendall et al. 2014, Ohlberger et al. 2018, Siegel et al. 2018, Ohlberger et al. 2023). Another caveat is that independent estimates of annual maturation rates separate from marine survival are unobtainable for wild populations because samples of subadults at sea (e.g., based on coded-wire tag recoveries) do not exist or are too small to be useful. While it would be ideal to model the distinct processes driving age at maturity and marine survival, what is feasible and perhaps more relevant is modeling adult age structure which implicitly includes both maturation and survival to adult return, as we have done.

Adult age and adult size in salmonids are not necessarily correlated over time (Kendall et al. 2014). Moreover, size and age are not always straightforwardly or monotonically related to reproductive output. Thus, a shift toward older age structure without a corresponding shift in size structure would likely yield the same population-level fecundity. Furthermore, given that the probability of survival to adulthood declines with increasing time spent at sea, it is possible for an older age structure to lead to lower overall reproductive output if the survival disadvantage outweighs the size-specific fecundity advantage. For example, Carvalho et al. (2023) found that increasing survival to maturity of age-4 and age-5 adult Sacramento River fall-run Chinook Salmon yielded greater spawner abundance, but delayed maturation decreased abundance. All else being equal, higher survival and larger, older spawners produce higher reproductive output, but this assumes the age-size correlation holds. This need not be the case for a specific population; alternatively, average age might increase while size-at-age declines, or average age and size might increase while survival declines. A joint investigation of the dynamics of survival and adult age structure would be a fruitful area for future research, especially to identify levers for management action that can increase the relative survival of older and larger spawners.

## Conclusion

Our analysis of Snake River spring/summer Chinook Salmon identified juvenile length and outmigration timing as the most important predictors of adult age at return. The link between juvenile size and age at maturity is well established, but the vital rates involved (i.e., age-specific marine survival and maturation) need further study. Outmigration timing is likely a proxy for multiple environmental conditions and processes experienced by juveniles (Gosselin et al. 2018). The environmental covariates we examined were less important predictors of adult age structure than anticipated, although this is consistent with previous studies of ocean conditions and age at maturity (Tattam et al. 2015). It is possible that more spatiotemporally or mechanistically refined indices of ocean conditions could reveal stronger relationships. Identifying predictors of maturation thresholds would likely entail mechanistic modeling of body size and growth in relation to physiological processes and environmental factors (Connolly et al. 2017). Our study points to the importance of developmental processes in the juvenile stage, and to a lesser extent river conditions during hydrosystem passage and ocean conditions during marine residence. Hatchery practices may have relatively direct impacts on juvenile growth and size, but habitat restoration is also clearly necessary for sustainable natural-origin populations. This highlights the importance of filling data and knowledge gaps in the effects of habitat restoration on early life history as it relates to adult age structure. Developmental and environmental effects on survival and age composition have potentially significant implications for population productivity and viability of endangered salmonids.

## Supporting information

Gosselin et al - SUPPL. MAT.

## Acknowledgements

Funding for this research was provided by the Bonneville Power Administration. We very much appreciate the field biologists who PIT-tagged the fish and made the data publicly available. We thank Kevin Malone for insightful discussions. We are grateful to Lisa Crozier, Michael Malick, Jan Ohlberger, and anonymous reviewers for their comments and help in improving the manuscript from earlier versions.

## Notes

https://github.com/Columbia-Basin-Research-CBR/sr-chinook-age-at-return

### Competing Interest Statement

The authors have declared no competing interest.

