## Supplementary material for "Biological, freshwater, and marine drivers of age at maturity in wild Chinook Salmon": Gosselin et al - SUPPL. MAT.

### Juvenile, freshwater, and marine covariates of age at maturity in returning adults of wild, spring/summer Chinook Salmon

Gosselin JL, Sandford BP, O'Brien CS, and Buhle ER

#### Supplemental Material

**Table S1.** Number of PIT-tagged, wild, spring/summer Snake River Basin Chinook Salmon returns by adult age.

| Origin | Adult return age | | | | All ages |
|---|---|---|---|---|---|
| | 3 (1-ocean) | 4 (2-ocean) | 5 (3-ocean) | 6 (4-ocean) | |
| Clearwater River | 4 | 86 | 36 | 0 | **126** |
| Grande Ronde/ Imnaha rivers | 77 | 729 | 210 | 1 | **1,017** |
| Lower Salmon River | 26 | 177 | 62 | 0 | **265** |
| Lower Snake River | 424 | 3,521 | 1,442 | 1 | **5,388** |
| Middle Fork Salmon River | 64 | 228 | 75 | 0 | **367** |
| South Fork Salmon River | 37 | 320 | 81 | 0 | **438** |
| Upper Salmon River | 29 | 313 | 76 | 0 | **418** |
| **All origin** | **661** | **5,374** | **1,982** | **2** | **8,019** |

**Table S2.** Number of PIT-tagged, wild, spring/summer Snake River Basin Chinook Salmon adult returns by juvenile tagging life stage and location and by adult age.

| Smolt migration year | All tag ages and locations | Tagging life stage & location and Adult return age | | | | | | | | | | | | | | |
|---|---|---|---|---|---|---|---|---|---|---|---|---|---|---|---|---|
| | | Parr&AboveLGR | | | | | Smolt&AboveLGR | | | | | Smolt&AtLGR | | | | |
| | | **3** | **4** | **5** | **6** | **all** | **3** | **4** | **5** | **6** | **all** | **3** | **4** | **5** | **6** | **all** |
| 1998 | 164 | 4 | 24 | 8 | 0 | **36** | 3 | 35 | 8 | 0 | **46** | 6 | 56 | 20 | 0 | **82** |
| 1999 | 469 | 1 | 20 | 5 | 0 | **26** | 4 | 66 | 20 | 0 | **90** | 14 | 285 | 54 | 0 | **353** |
| 2000 | 1103 | 1 | 12 | 12 | 0 | **25** | 2 | 26 | 46 | 0 | **74** | 26 | 394 | 583 | 1 | **1,004** |
| 2001 | 191 | 0 | 1 | 1 | 0 | **2** | 0 | 9 | 3 | 0 | **12** | 21 | 125 | 31 | 0 | **177** |
| 2002 | 445 | 1 | 6 | 2 | 0 | **9** | 1 | 9 | 4 | 0 | **14** | 31 | 310 | 81 | 0 | **422** |
| 2003 | 122 | 0 | 6 | 1 | 0 | **7** | 1 | 9 | 5 | 0 | **15** | 2 | 66 | 32 | 0 | **100** |
| 2004 | 138 | 0 | 9 | 10 | 0 | **19** | 1 | 29 | 19 | 1 | **50** | 3 | 49 | 17 | 0 | **69** |
| 2005 | 85 | 0 | 15 | 3 | 0 | **18** | 1 | 12 | 7 | 0 | **20** | 1 | 30 | 16 | 0 | **47** |
| 2006 | 318 | 3 | 32 | 8 | 0 | **43** | 0 | 24 | 10 | 0 | **34** | 11 | 186 | 44 | 0 | **241** |
| 2007 | 390 | 1 | 34 | 5 | 0 | **40** | 6 | 45 | 16 | 0 | **67** | 29 | 222 | 32 | 0 | **283** |
| 2008 | 1212 | 28 | 195 | 59 | 0 | **282** | 7 | 85 | 47 | 0 | **139** | 91 | 529 | 171 | 0 | **791** |
| 2009 | 749 | 6 | 99 | 34 | 0 | **139** | 14 | 123 | 57 | 0 | **194** | 20 | 279 | 117 | 0 | **416** |
| 2010 | 344 | 19 | 41 | 18 | 0 | **78** | 17 | 52 | 23 | 0 | **92** | 25 | 110 | 39 | 0 | **174** |
| 2011 | 184 | 3 | 43 | 7 | 0 | **53** | 1 | 40 | 8 | 0 | **49** | 7 | 59 | 16 | 0 | **82** |
| 2012 | 569 | 29 | 114 | 10 | 0 | **153** | 21 | 101 | 13 | 0 | **135** | 53 | 197 | 31 | 0 | **281** |
| 2013 | 500 | 7 | 53 | 19 | 0 | **79** | 10 | 84 | 26 | 0 | **120** | 26 | 211 | 64 | 0 | **301** |
| 2014 | 226 | 4 | 35 | 9 | 0 | **48** | 5 | 54 | 15 | 0 | **74** | 10 | 77 | 17 | 0 | **104** |
| 2015 | 38 | 0 | 11 | 0 | 0 | **11** | 2 | 8 | 1 | 0 | **11** | 4 | 10 | 2 | 0 | **16** |
| 2016 | 140 | 7 | 18 | 4 | 0 | **29** | 3 | 38 | 4 | 0 | **45** | 9 | 54 | 3 | 0 | **66** |
| 2017 | 72 | 0 | 9 | 5 | 0 | **14** | 1 | 10 | 1 | 0 | **12** | 4 | 38 | 4 | 0 | **46** |
| 2018 | 207 | 3 | 28 | 0 | 0 | **31** | 2 | 38 | 4 | 0 | **44** | 13 | 106 | 13 | 0 | **132** |
| 2019 | 120 | 4 | 28 | 7 | 0 | **39** | 2 | 15 | 3 | 0 | **20** | 3 | 51 | 7 | 0 | **61** |
| 2020 | 233 | 17 | 113 | 13 | 0 | **143** | 10 | 72 | 8 | 0 | **90** | 0 | 0 | 0 | 0 | **0** |
| **All years** | **8019** | **138** | **946** | **240** | **0** | **1324** | **114** | **984** | **348** | **1** | **1447** | **409** | **3444** | **1394** | **1** | **5248** |

**Table S3.** Number of PIT-tagged, wild, spring/summer Snake River Basin Chinook Salmon adult returns by passage type and adult age.

| Smolt migration year | Passage type and adult return age | | | | | | | | | |
|---|---|---|---|---|---|---|---|---|---|---|
| | In-river | | | | | Transported | | | | |
| | 3 (1-ocean) | 4 (2-ocean) | 5 (3-ocean) | 6 (4-ocean) | All | 3 (1-ocean) | 4 (2-ocean) | 5 (3-ocean) | 6 (4-ocean) | All |
| 1998 | 8 | 92 | 28 | 0 | **128** | 5 | 23 | 8 | 0 | **36** |
| 1999 | 7 | 192 | 48 | 0 | **247** | 12 | 179 | 31 | 0 | **222** |
| 2000 | 21 | 295 | 484 | 0 | **800** | 8 | 137 | 157 | 1 | **303** |
| 2001 | 0 | 10 | 3 | 0 | **13** | 21 | 125 | 32 | 0 | **178** |
| 2002 | 10 | 129 | 29 | 0 | **168** | 23 | 196 | 58 | 0 | **277** |
| 2003 | 2 | 35 | 12 | 0 | **49** | 1 | 46 | 26 | 0 | **73** |
| 2004 | 1 | 19 | 7 | 1 | **28** | 3 | 68 | 39 | 0 | **110** |
| 2005 | 1 | 19 | 9 | 0 | **29** | 1 | 38 | 17 | 0 | **56** |
| 2006 | 5 | 91 | 25 | 0 | **121** | 9 | 151 | 37 | 0 | **197** |
| 2007 | 14 | 125 | 28 | 0 | **167** | 22 | 176 | 25 | 0 | **223** |
| 2008 | 31 | 276 | 92 | 0 | **399** | 95 | 533 | 185 | 0 | **813** |
| 2009 | 14 | 233 | 89 | 0 | **336** | 26 | 268 | 119 | 0 | **413** |
| 2010 | 27 | 90 | 29 | 0 | **146** | 34 | 113 | 51 | 0 | **198** |
| 2011 | 6 | 90 | 23 | 0 | **119** | 5 | 52 | 8 | 0 | **65** |
| 2012 | 71 | 296 | 35 | 0 | **402** | 32 | 116 | 19 | 0 | **167** |
| 2013 | 19 | 129 | 40 | 0 | **188** | 24 | 219 | 69 | 0 | **312** |
| 2014 | 13 | 112 | 24 | 0 | **149** | 6 | 54 | 17 | 0 | **77** |
| 2015 | 1 | 12 | 1 | 0 | **14** | 5 | 17 | 2 | 0 | **24** |
| 2016 | 15 | 64 | 9 | 0 | **88** | 4 | 46 | 2 | 0 | **52** |
| 2017 | 2 | 43 | 8 | 0 | **53** | 3 | 14 | 2 | 0 | **19** |
| 2018 | 5 | 79 | 2 | 0 | **86** | 13 | 93 | 15 | 0 | **121** |
| 2019 | 6 | 40 | 11 | 0 | **57** | 3 | 54 | 6 | 0 | **63** |
| 2020 | 22 | 144 | 19 | 0 | **185** | 5 | 41 | 2 | 0 | **48** |
| **All** | **301** | **2,615** | **1,055** | **1** | **3,972** | **360** | **2,759** | **927** | **1** | **4,047** |

**Table S4.** Estimated regression coefficients (β) for covariate effects on age at maturity of wild, spring/summer Snake River Chinook Salmon adult returns in the model that includes DOY at LGR (i.e., model in main paper). The estimated error and the lower and upper confidence limits (CL) are included.

| $\tau$ intercepts and β effects | Estimate | Estimated Error | Lower 90% CL | Upper 90% CL |
|---|---|---|---|---|
| $\tau_1$ | -1.35 | 0.12 | -1.55 | -1.16 |
| $\tau_2$ | 0.92 | 0.12 | 0.74 | 1.11 |
| Length | -0.18 | 0.04 | -0.25 | -0.12 |
| Smolt&AboveLGR | 0.25 | 0.09 | 0.11 | 0.39 |
| Smolt&AtLGR | 0.12 | 0.13 | -0.10 | 0.34 |
| Origin Lower Snake | 0.16 | 0.11 | -0.02 | 0.35 |
| Origin Clearwater | 0.18 | 0.12 | -0.01 | 0.37 |
| Origin SF Salmon | -0.17 | 0.08 | -0.30 | -0.03 |
| Origin Lower Salmon | 0.04 | 0.09 | -0.10 | 0.18 |
| Origin MF Salmon | -0.34 | 0.09 | -0.48 | -0.20 |
| Origin Upper Salmon | 0.08 | 0.08 | -0.05 | 0.21 |
| LGR passage day of year | 0.21 | 0.02 | 0.18 | 0.24 |
| LGR flow | 0.03 | 0.02 | 0.00 | 0.07 |
| Transported hydrosystem passage | -0.11 | 0.03 | -0.16 | -0.06 |
| NPGO index | 0.14 | 0.06 | 0.03 | 0.24 |
| Length x Smolt&AboveLGR | 0.08 | 0.05 | 0.00 | 0.17 |
| Length x Smolt&AtLGR | -0.05 | 0.05 | -0.14 | 0.04 |

**Table S5.** Estimated regression coefficients (β) for covariate effects on age of wild, spring/summer Snake River Chinook Salmon adult returns in the model that includes river temperature at LGR in place of day-of-year of LGR passage. The estimated error and the lower and upper confidence limits (CL) are included.

| $\tau$ intercepts and β effects | Estimate | Estimated Error | Lower 90% CL | Upper 90% CL |
|---|---|---|---|---|
| $\tau_1$ | -1.29 | 0.11 | -1.47 | -1.10 |
| $\tau_2$ | 0.98 | 0.11 | 0.80 | 1.17 |
| Length | -0.19 | 0.04 | -0.25 | -0.12 |
| Smolt&AboveLGR | 0.28 | 0.09 | 0.14 | 0.42 |
| Smolt&AtLGR | 0.15 | 0.14 | -0.07 | 0.37 |
| Origin Lower Snake | 0.17 | 0.11 | -0.01 | 0.36 |
| Origin Clearwater | 0.20 | 0.11 | 0.01 | 0.39 |
| Origin SF Salmon | -0.16 | 0.08 | -0.29 | -0.02 |
| Origin Lower Salmon | 0.02 | 0.09 | -0.12 | 0.16 |
| Origin MF Salmon | -0.34 | 0.09 | -0.48 | -0.19 |
| Origin Upper Salmon | 0.10 | 0.08 | -0.03 | 0.23 |
| LGR temperature | 0.18 | 0.02 | 0.15 | 0.21 |
| LGR flow | 0.07 | 0.02 | 0.04 | 0.11 |
| Transported hydrosystem passage | -0.10 | 0.03 | -0.15 | -0.05 |
| NPGO index | 0.16 | 0.06 | 0.06 | 0.26 |
| Length x Smolt&AboveLGR | 0.07 | 0.05 | -0.01 | 0.16 |
| Length x Smolt&AtLGR | -0.05 | 0.05 | -0.14 | 0.04 |

**Table S6.** Estimated regression coefficients (β) for covariate effects on age of wild, spring/summer Snake River Chinook Salmon adult returns in the model that includes only biological / behavioural covariates. The estimated error and the lower and upper confidence limits (CL) are included.

| $\tau$ intercepts and β effects | Estimate | Estimated Error | Lower 90% CL | Upper 90% CL |
|---|---|---|---|---|
| $\tau_1$ | -1.24 | 0.11 | -1.43 | -1.05 |
| $\tau_2$ | 1.03 | 0.11 | 0.84 | 1.22 |
| Length | -0.19 | 0.04 | -0.25 | -0.12 |
| Smolt&AboveLGR | 0.24 | 0.09 | 0.10 | 0.38 |
| Smolt&AtLGR | 0.10 | 0.14 | -0.13 | 0.32 |
| Origin Lower Snake | 0.16 | 0.11 | -0.03 | 0.34 |
| Origin Clearwater | 0.19 | 0.12 | -0.01 | 0.38 |
| Origin SF Salmon | -0.16 | 0.08 | -0.29 | -0.03 |
| Origin Lower Salmon | 0.04 | 0.08 | -0.10 | 0.18 |
| Origin MF Salmon | -0.36 | 0.09 | -0.50 | -0.22 |
| Origin Upper Salmon | 0.08 | 0.08 | -0.05 | 0.20 |
| LGR passage day of year | 0.22 | 0.02 | 0.19 | 0.24 |
| Length x Smolt&AboveLGR | 0.08 | 0.05 | -0.01 | 0.17 |
| Length x Smolt&AtLGR | -0.06 | 0.05 | -0.15 | 0.03 |

**Table S7.** Estimated regression coefficients (β) for covariate effects on age of wild, spring/summer Snake River Chinook Salmon adult returns in the model that includes only freshwater covariates

| $\tau$ intercepts and β effects | Estimate | Estimated Error | Lower 90% CL | Upper 90% CL |
|---|---|---|---|---|
| $\tau_1$ | -1.44 | 0.08 | -1.58 | -1.31 |
| $\tau_2$ | 0.81 | 0.08 | 0.68 | 0.94 |
| LGR temperature | 0.17 | 0.02 | 0.14 | 0.20 |
| LGR flow | 0.09 | 0.02 | 0.05 | 0.12 |
| Transported hydrosystem passage | -0.12 | 0.03 | -0.17 | -0.08 |

**Table S8.** Estimated regression coefficients (β) for covariate effects on age of wild, spring/summer Snake River Chinook Salmon adult returns in the model that includes freshwater covariates, but not river temperature.

| $\tau$ intercepts and β effects | Estimate | Estimated Error | Lower 90% CL | Upper 90% CL |
|---|---|---|---|---|
| $\tau_1$ | -1.44 | 0.09 | -1.58 | -1.29 |
| $\tau_2$ | 0.79 | 0.09 | 0.65 | 0.94 |
| LGR river flow | 0.18 | 0.02 | 0.15 | 0.21 |
| Transported hydrosystem passage | -0.07 | 0.03 | -0.12 | -0.02 |

**Table S9.** Estimated regression coefficients (β) for covariate effects on age of wild, spring/summer Snake River Chinook Salmon adult returns in the model that includes only the marine covariate.

| $\tau$ intercepts and β effects | Estimate | Estimated Error | Lower 90% CL | Upper 90% CL |
|---|---|---|---|---|
| $\tau_1$ | -1.44 | 0.08 | -1.57 | -1.31 |
| $\tau_2$ | 0.77 | 0.08 | 0.64 | 0.89 |
| NPGO index | 0.12 | 0.06 | 0.01 | 0.22 |

**Table S10.** Leave-one-out cross-validation (LOO) of models examined.

| Model run name and referenced output | elpd_diff | elpd_se_diff | looic | se_looic |
|---|---|---|---|---|
| fit_DOY_npgo3 (Table S4) | 0.0 | 0.0 | 12013.2 | 124.4 |
| fit_fish (Table S6) | -5.6 | 3.9 | 12024.6 | 124.3 |
| fit_TEMP_npgo3 (Table S5) | -20.4 | 6.1 | 12054.0 | 124.4 |
| fit_freshwater1 (Table S7) | -57.1 | 11.6 | 12127.5 | 124.8 |
| fit_freshwater2 (Table S8) | -100.4 | 14.9 | 12213.9 | 124.7 |
| fit_marine (Table S9) | -150.9 | 17.9 | 12315.1 | 124.7 |

**Table S11.** Estimated random effects error ($e$) associated with smolt migration year $t$ in the probit regression model, Eq. 1.

| Year | Mean | 90% CI |
|---|---|---|
| 1998 | 0.0819 | (-0.1566, 0.3187) |
| 1999 | -0.2031 | (-0.4347, 0.029) |
| 2000 | 0.8038 | (0.5825, 1.0294) |
| 2001 | -0.1839 | (-0.3963, 0.0258) |
| 2002 | -0.1511 | (-0.3057, 0.0003) |
| 2003 | 0.2965 | (0.0838, 0.509) |
| 2004 | 0.5233 | (0.3079, 0.7429) |
| 2005 | 0.4061 | (0.1768, 0.6399) |
| 2006 | -0.0526 | (-0.2244, 0.1171) |
| 2007 | -0.2237 | (-0.4053, -0.043) |
| 2008 | -0.1243 | (-0.2865, 0.0373) |
| 2009 | 0.112 | (-0.0469, 0.2737) |
| 2010 | -0.3333 | (-0.5093, -0.156) |
| 2011 | -0.1096 | (-0.2947, 0.0747) |
| 2012 | -0.4911 | (-0.641, -0.3447) |
| 2013 | 0.0382 | (-0.1206, 0.1943) |
| 2014 | 0.0542 | (-0.1192, 0.2291) |
| 2015 | -0.162 | (-0.4692, 0.141) |
| 2016 | -0.2478 | (-0.4775, -0.0194) |
| 2017 | 0.0942 | (-0.2024, 0.3926) |
| 2018 | -0.053 | (-0.3049, 0.1926) |
| 2019 | 0.1047 | (-0.1493, 0.3645) |
| 2020 | -0.1286 | (-0.3615, 0.1003) |

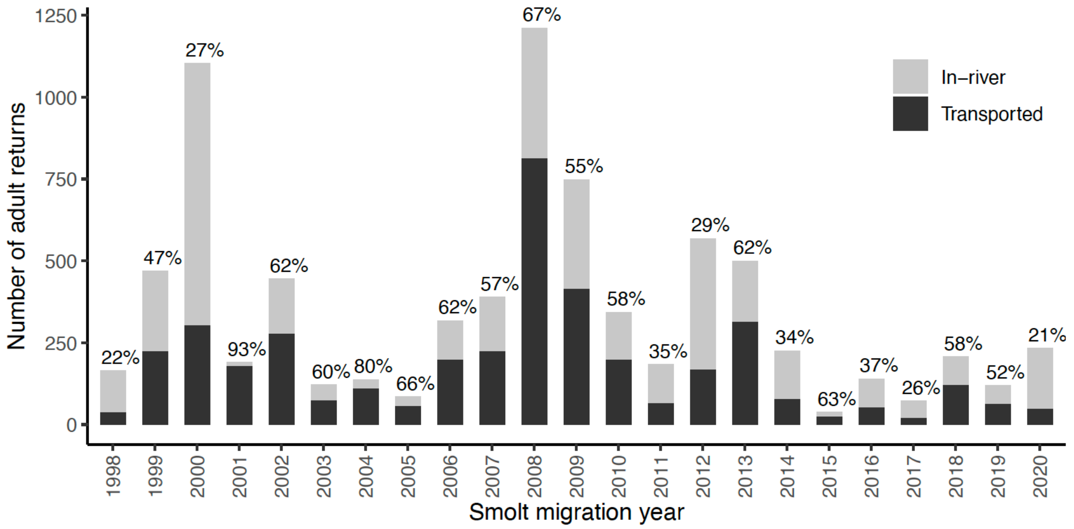


**Figure S1**. Number of PIT-tagged wild, spring/summer, Snake River Chinook Salmon adult returns by smolt migration year and subdivided by juvenile passage type (in-river or transported). Percentage of juveniles transported each year are reported above each stacked bar.

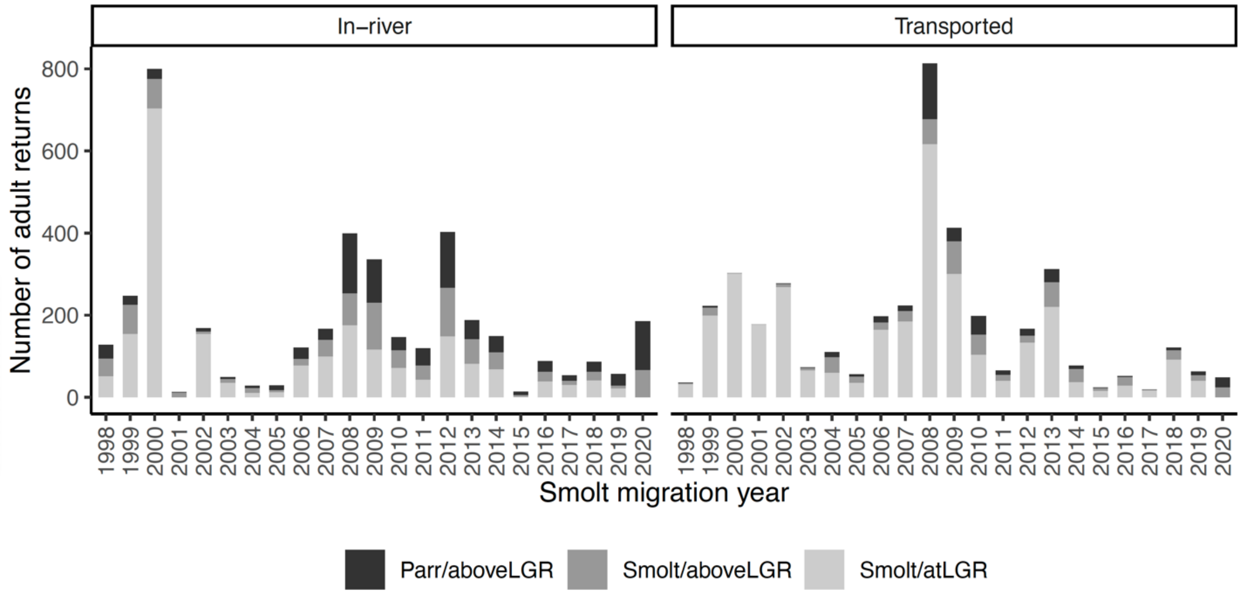


**Figure S2.** Number of wild, spring/summer, Snake River Chinook Salmon adult returns by smolt migration year, subdivided by passage type and life stage / location at time of tagging.

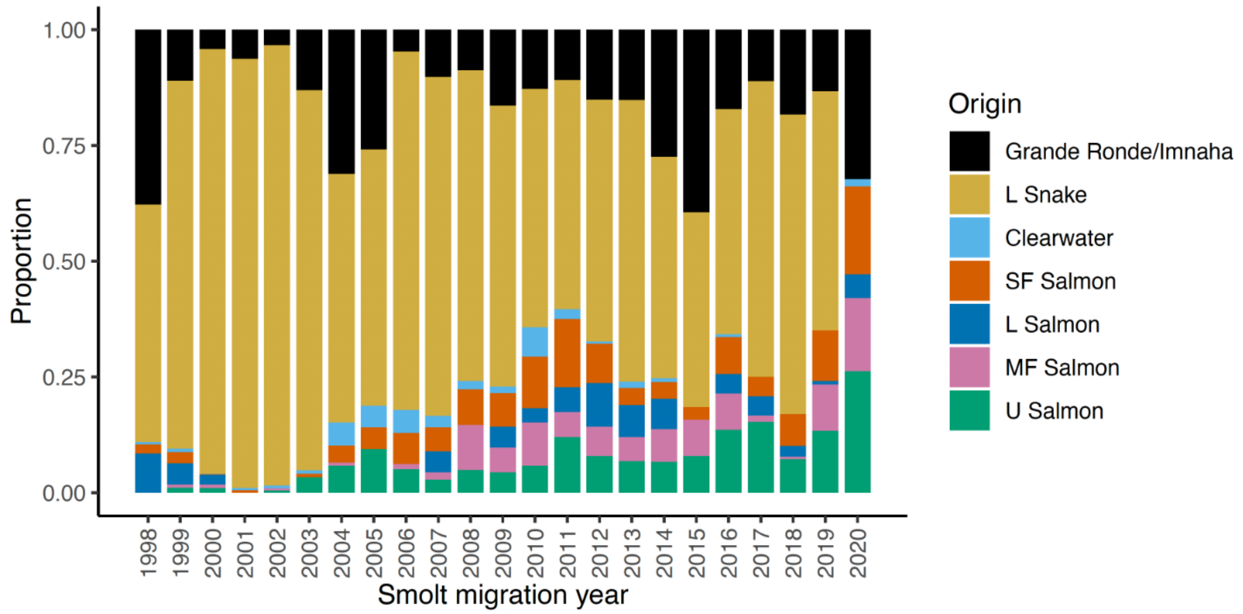


**Figure S3.** Proportion of adult returns by smolt migration year by origin of wild, spring/summer, Snake River Chinook Salmon adult returns tagging and release.

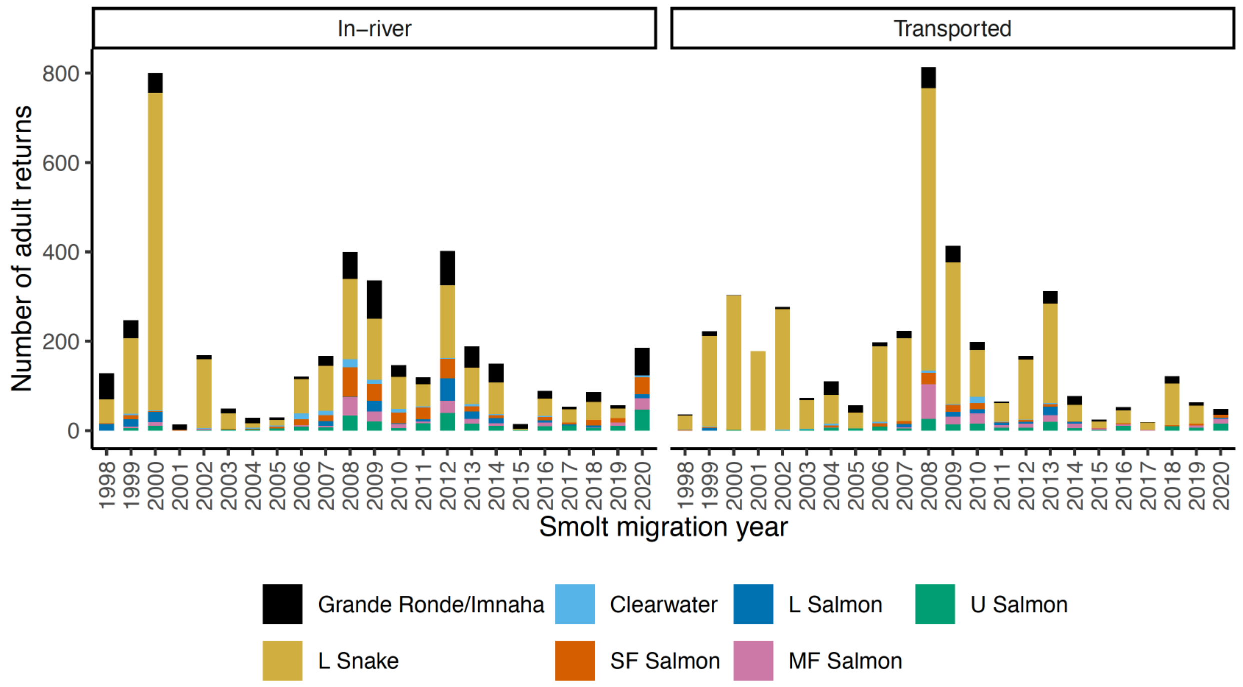


**Figure S4.** Number of adult returns by smolt migration year, subdivided by passage type and origin of wild, spring/summer, Snake River Chinook Salmon adult returns tagging and release.

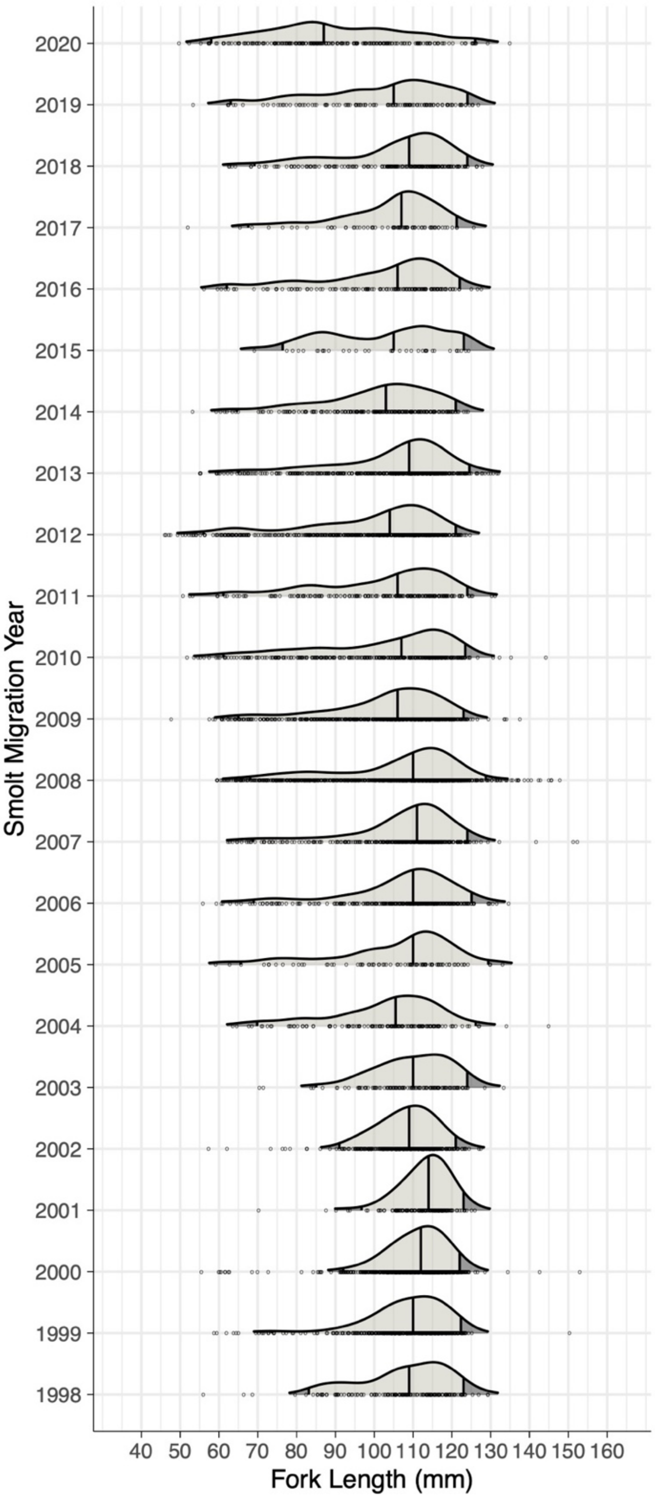


**Figure S5.** Fork length of parr and smolt wild, spring/summer Snake River Chinook Salmon measured at time of tagging and release, above or at Lower Granite Dam, by smolt migration year. Density curves with a relative minimum height of 0.03 are shown, along with light grey area representing the 95% CI. Vertical lines under the density curves, from left to right, respectively represent 2.5$^{th}$, 50$^{th}$, and 97.5$^{th}$ percentile.

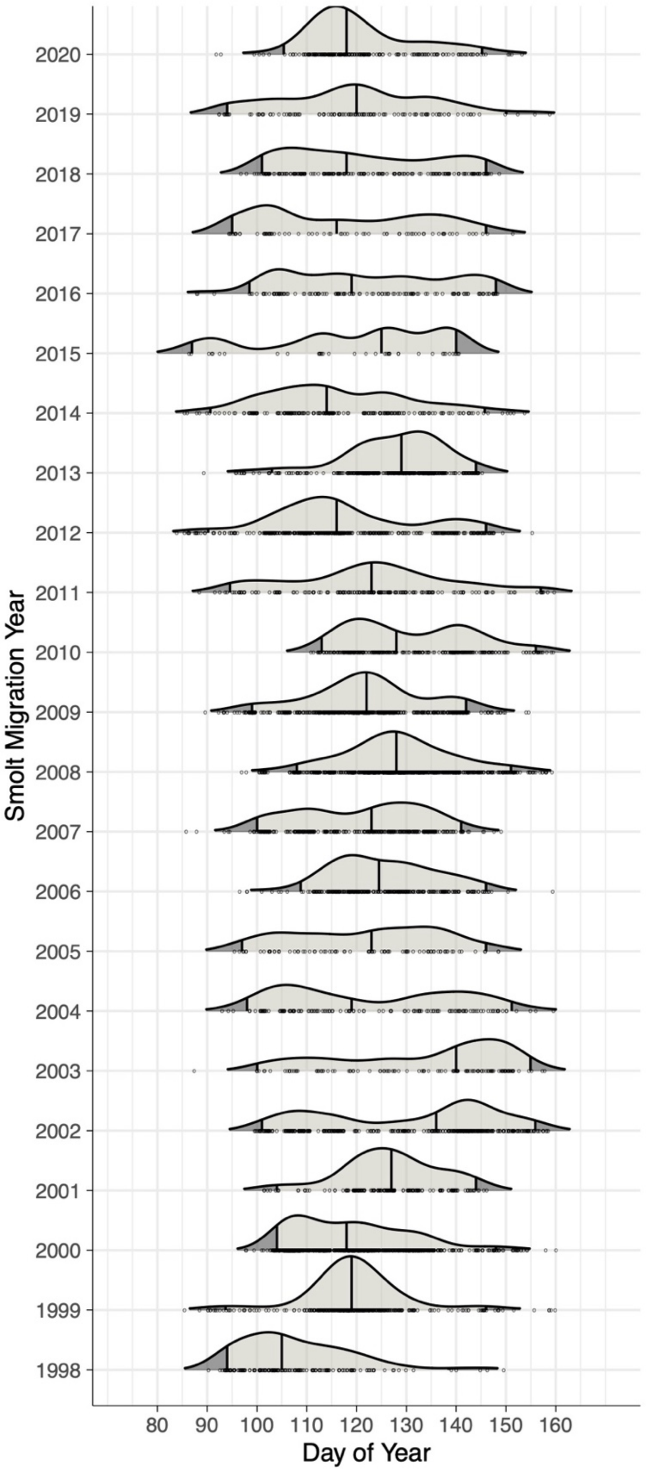


**Figure S6.** Lower Granite Dam passage day-of-year in each smolt migration year for wild, spring/summer, Snake River Chinook Salmon. Density curves with a relative minimum height of 0.03 are shown, along with light grey area representing the 95% CI. Vertical lines under the density curves, from left to right, respectively represent $2.5^{th}$, $50^{th}$, and $97.5^{th}$ percentiles.

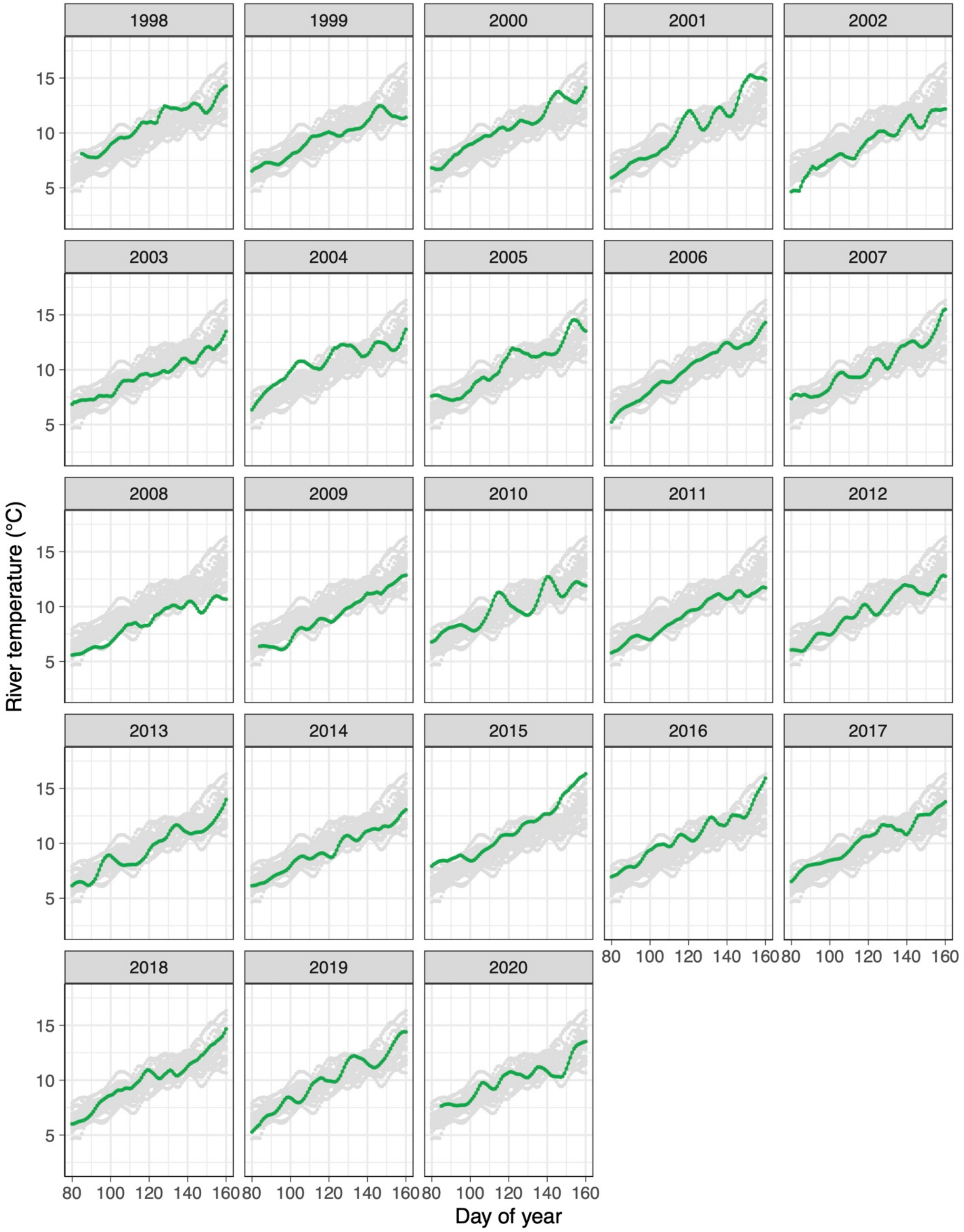


Figure S7. Lower Granite Dam river temperature in all years examined (grey) and in smolt migration year (green).

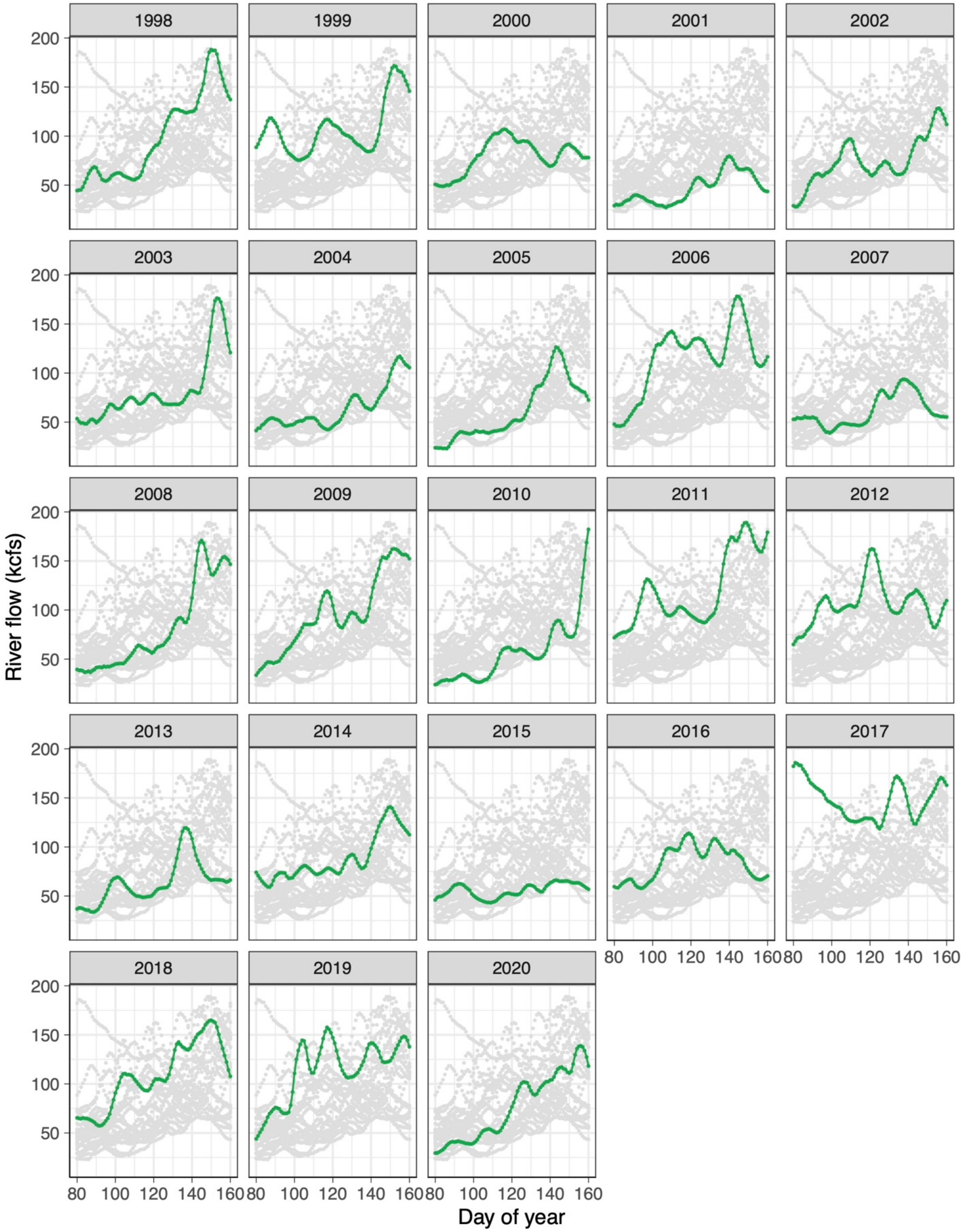


**Figure S8.** Lower Granite Dam river flow in all years examined (grey) and in smolt migration year (green).

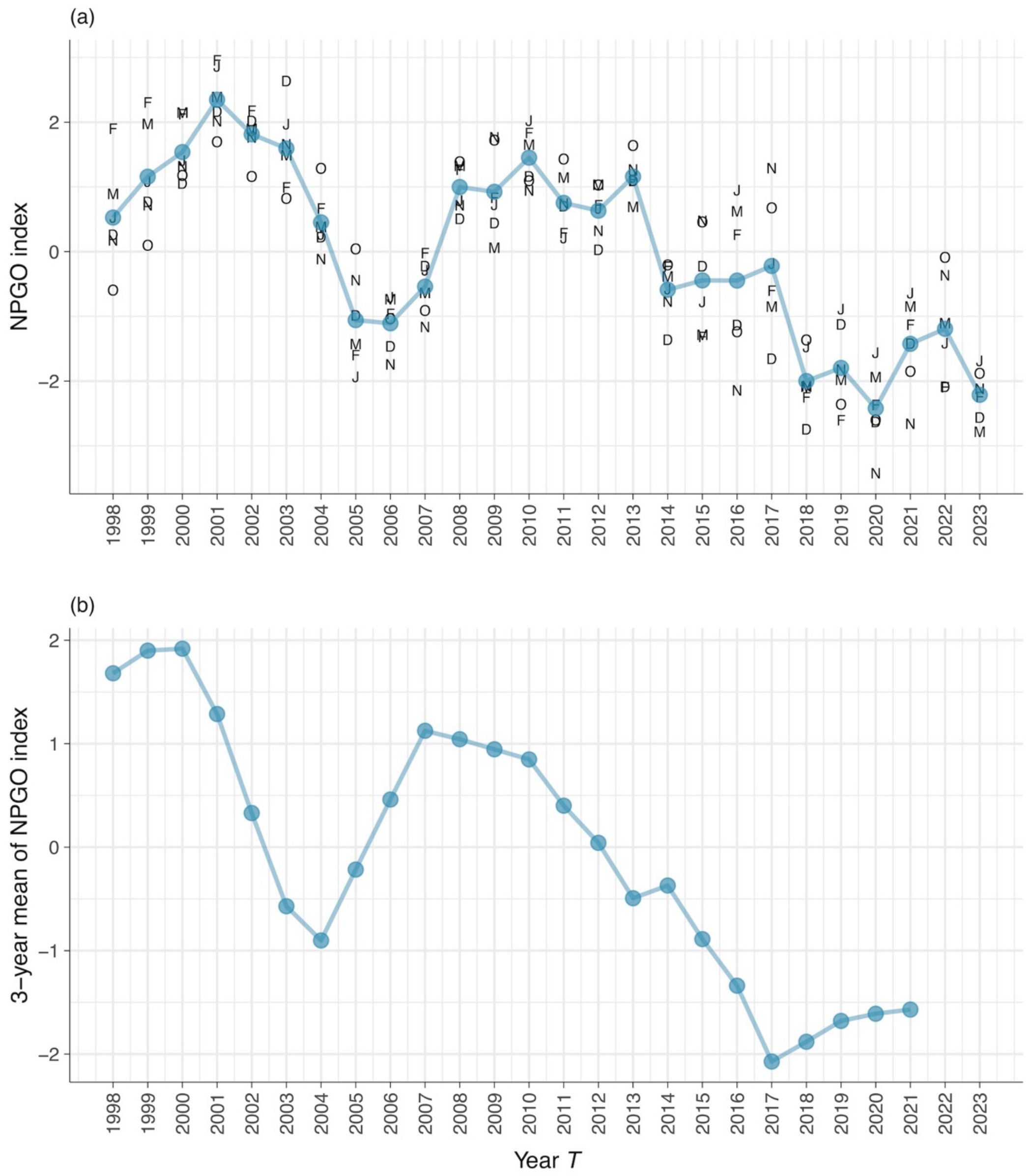


**Figure S9.** North Pacific Gyre Oscillation (NPGO) index (a) averaged over October in year *T-1* through March in year *T*, and (b) as a 3-year mean of the NPGO index in (a) starting with year *T* on the x-axis.
